# Microglia shape hippocampal networks but are dispensable for pruning of synapses during development

**DOI:** 10.1101/2023.10.05.560092

**Authors:** Michael Surala, Luna Soso-Zdravkovic, David Munro, Ali Rifat, Koliane Ouk, Imre Vida, Josef Priller, Christian Madry

## Abstract

Microglia, the brain-resident macrophages, are believed to sculpt developing neural circuits by eliminating excess synapses in a process called synaptic pruning, removing apoptotic neurons, and promoting neuronal survival. To elucidate the role of microglia during embryonic and postnatal brain development, we used a mouse model deficient in microglia throughout life as a consequence of deletion of the fms-intronic regulatory element (FIRE) in the *Csf1r* locus. Surprisingly, young adult *Csf1r*^ΔFIRE/ΔFIRE^ mice displayed no changes in excitatory synapse number and spine density of CA1 hippocampal neurons compared to *Csf1r*^+/+^ littermates. However, CA1 neurons were less excitable, received less CA3 Schaffer collateral-mediated excitatory input and showed altered synaptic properties, but this did not affect novel object recognition. Cytokine profiling indicated an anti-inflammatory state along with increases in ApoE levels and astrocyte reactivity in brains of *Csf1r*^ΔFIRE/ΔFIRE^ mice. Notably, these morphological and functional changes in *Csf1r*^ΔFIRE/ΔFIRE^ mice closely resemble the effects of acute microglial depletion in adult mice that had undergone normal development. Thus, our findings challenge the prevailing view that microglia are indispensable for the establishment of neural networks during development.

## Introduction

Microglia are the primary innate immune cells in the central nervous system (CNS). Beyond fulfilling classical immunological roles to protect the brain from injury or invading pathogens, microglia are increasingly recognized as key players involved in shaping neural function both in the developing and adult CNS (Thion and Garel, 2017, Pósfai et al., 2019). At early embryonic stages, when other glial populations such as astrocytes have not yet been generated, yolk sac-derived erythromyeloid progenitors engraft in the brain, differentiate into microglia, proliferate and rapidly colonize all brain areas until reaching maturity and steady-state cell densities early postnatally (Prinz and Priller, 2014). Thereafter, they reside as long-lived tissue macrophages in the brain parenchyma and, under physiological conditions, maintain their population through self-renewal (Askew and Gomez-Nicola, 2018).

Since microglia arise around the same time as neurons in the developing CNS, they are in an ideal position to interact with them from early on. Indeed, microglia have been shown to promote embryonic and adult neurogenesis (Vukovic et al., 2012, Xavier et al., 2015), as well as neuronal survival (Morgan et al., 2004, Ueno et al., 2013). They regulate axon guidance and the positioning of neocortical interneurons (Squarzoni et al., 2014), contribute to the migration and differentiation of neural precursor cells (Aarum et al., 2003, Antony et al., 2011), and eliminate newborn neurons and astrocytes by phagocytosis (Sierra et al., 2010, Cunningham et al., 2013, VanRyzin et al., 2019, Diaz-Aparicio et al., 2020). In late embryonic and early postnatal development, newly generated neurons undergo extensive synaptogenesis, followed by a period of net synapse elimination in a process called “synaptic pruning”, which is critical for the maturation and refinement of neural circuits. Microglia participate in this process by eliminating unwanted or excess synapses of both excitatory and inhibitory circuits involving microglial-specific receptor pathways such as the fractalkine receptor (CX3CR1), the triggering receptor expressed on myeloid cells 2 (TREM2), or complement receptor CR3 (Paolicelli et al., 2011, Schafer et al., 2012, Zhan et al., 2014, Filipello et al., 2018, Gunner et al., 2019, Favuzzi et al., 2021). Apart from the elimination of synapses, microglia may also induce their formation and presynaptic rearrangement (Trapp et al., 2007, Miyamoto et al., 2016, Weinhard et al., 2018). Importantly, on the functional level, microglia regulate synaptic transmission and plasticity, thus affecting network function and consequently memory formation and behavior (Paolicelli et al., 2011, Parkhurst et al., 2013, Zhan et al., 2014, Basilico et al., 2019, Merlini et al., 2021, Wang et al., 2020, Basilico et al., 2022). Microglial interactions with neurons occur at specific cell-cell contacts (Cserép et al., 2020), and depend on neuronal activity (Tremblay et al., 2010, Li et al., 2012, Liu et al., 2019; Stowell et al., 2019; Cserép et al., 2020, Badimon et al., 2020). Remarkably, by reducing the discharge rate of excessively active neurons, microglia reinstate homeostasis by protecting neurons from adopting potentially harmful hyperactive states (Li et al., 2012, Cserép et al., 2020, Badimon et al., 2020).

Collectively, the existing literature ascribes to microglia a crucial role in CNS development and function, leading to the hypothesis that without the involvement of microglia these processes may be significantly impaired with deficits in neuronal maturation and function on the cellular, synaptic and network level. This view is further supported by recent studies in adult mice in which microglia have been transiently depleted, resulting in changes in both glutamatergic and GABAergic transmission (Liu et al., 2021, Ma et al., 2020, Basilico et al., 2022, Du et al., 2022).

Despite increasing knowledge of microglia as active sculptors of neural circuits, their global role in shaping brain function, including the contribution of other microglial-independent mechanisms, still remains largely unclear. This is mainly due to the fact that existing studies employed models with functionally impaired or transiently depleted microglia at postnatal stages, so that microglia were either always present, albeit functionally compromised, to maintain interactions with other cells, or they were only absent from later postnatal stages onward.

The proliferation, differentiation and survival of cells of the mononuclear phagocyte system depends upon signals from the macrophage colony-stimulating factor (CSF1) receptor (CSF1R) which is expressed exclusively in cells of this lineage. Dominant and recessive mutations in the CSF1R gene in zebrafish, mice, rats and humans are associated with the loss of microglia and most tissue macrophage populations, with complex impacts on postnatal development (Hume et al., 2020). In the rat, complete loss of microglia and brain-associated macrophages as a consequence of homozygous mutation of the *Csf1r* locus has remarkably little impact on pre- and postnatal brain development (Keshvari et al., 2021; Patkar et al., 2021), and in C57BL/6J mice, the perinatal lethality of *Csf1r* mutation can be overcome by neonatal transfer of wild-type bone marrow cells (Bennett et al., 2018). These findings suggest that developmental functions of microglia may be partly redundant.

In this study, we sought to elucidate the influence of microglia on shaping neuronal properties during brain development by using a mouse model which is entirely deficient in microglia throughout all stages of life, including embryonal development (Rojo et al., 2019). This line was generated by germ-line deletion of the *fms*-intronic regulatory element (FIRE) in the *Csf1r* locus, which is required for expression of the CSF1 receptor in bone marrow progenitors and blood monocytes. At least on a mixed genetic background, *Csf1r*^ΔFIRE/ΔFIRE^ mice are healthy, fertile and develop normally without apparent behavioral abnormalities but lack microglia and subsets of brain-associated macrophages, as well as macrophages in heart, skin, kidney and peritoneal cavity (Rojo et al., 2019; McNamara et al., 2023). The absence of microglia in this line had little effect on gene expression in the hippocampus, aside from the loss of the microglial gene signature (Rojo et al., 2019).

We characterized the effect of the developmental absence of microglia in *Csf1r*^ΔFIRE/ΔFIRE^ mice on the glutamatergic network in the hippocampus in young adult mice, at a stage when synaptic pruning has been largely completed (Jawaid et al., 2018). Specifically, we examined the morphological and functional properties of hippocampal pyramidal neurons on the cellular, synaptic and microcircuit level by whole-cell patch-clamp electrophysiology. The lack of microglia did not affect excitatory synapse number nor spine density and morphology of CA1 neurons but resulted in a weakened glutamatergic transmission in *Csf1r*^ΔFIRE/ΔFIRE^ mice compared to littermate controls. We further investigated the consequences of microglial deficiency on nonspatial memory in the novel object recognition test which involves the hippocampus, and on the immunological milieu in brains of *Csf1r*^ΔFIRE/ΔFIRE^ mice by biochemical and ELISA-based analyses. Our data show that object memory was not affected and cytokine levels were only mildly altered towards an anti-inflammatory state, while ApoE levels were increased and astrocytes were reactive in *Csf1r*^ΔFIRE/ΔFIRE^ mice. In summary, we found little evidence of an absolute requirement for microglia in normal neural development under healthy conditions.

## Results

### Absence of microglia has no effect on CA1 excitatory synapse number and spine density and morphology

Morphological properties of hippocampal CA1 pyramidal neurons were determined in acute brain slices from young adult 6-10-week-old *Csf1r*^ΔFIRE/ΔFIRE^ mice and controls at post-pruning stages (Jawaid et al., 2018). In line with previous findings (Rojo et al., 2019), we observed a complete absence of parenchymal Iba1 immunoreactivity in hippocampal and neocortical regions of *Csf1r*^ΔFIRE/ΔFIRE^ mice compared to wild-type (WT, *Csf1r*^+/+^) littermates (Fig 1A). We chose the hippocampus as a particularly well characterized brain region for microglia-neuron interactions (Paolicelli et al., 2011, Parkhurst et al., 2013, Zhan et al., 2014, Basilico et al., 2020), which would allow us to compare our findings with recent studies investigating the neuronal consequences of transient depletion of microglia in adult WT mice (Parkhurst et al., 2013, Basilico et al., 2022, Du et al., 2022).

**Figure 1.**
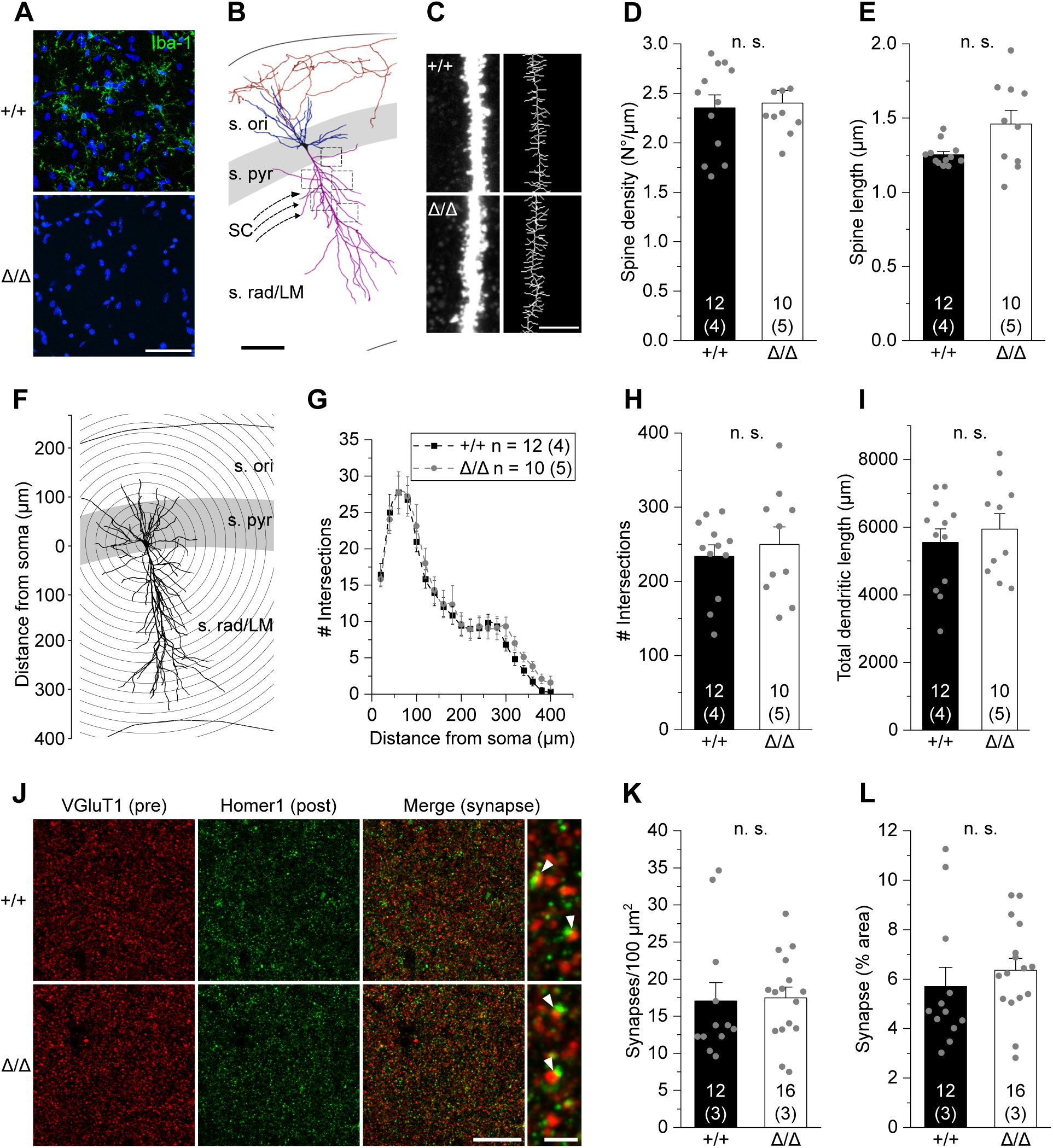
Absence of microglia has no effect on CA1 excitatory synapse number, spine density and morphology. A. Specimen confocal images illustrating the lack of microglia by Iba1 immunoreactivity (green) in acute hippocampal slices of *Csf1r*^ΔFIRE/ΔFIRE^ (Δ/Δ) compared to WT littermates (+/+). DAPI labeling of cellular nuclei in blue. Scale bar, 50 µm. B. 3D-reconstruction of a biocytin-filled CA1 pyramidal cell (black, cell body; blue, basal dendrite; magenta, apical dendrite; red, axon), depicting the excitatory input via the Schaffer collaterals (SC) and hippocampal layers. (s. ori – *stratum oriens*, s. pyr – *stratum pyramidale*, s. rad/LM – *stratum radiatum/lacunosum-moleculare*). Squared boxes indicate five different apical regions of interest for spine analysis. Individual values were averaged to obtain a grand average per cell. Scale bar, 50 µm. C. Representative segments of apical dendrites at high power (left) from which skeletonized branches were generated (right). Scale bar, 5 µm. D, E. Analysis of mean apical spine density (D) and spine length (E). F. Schematic representation of Sholl analysis of a biocytin-filled CA1 pyramidal cell. Number of intersections between dendrites and concentric spheres centred around the soma was determined at increasing distances with 20 μm increments. G-I. Sholl analysis-derived values of the number of intersections with Sholl radii at increasing distance from the soma (G) and resulting total number of intersections (H) and total dendritic length (I) of CA1 pyramidal cells. J. Confocal images showing VGluT1-labeled presynaptic puncta (red) and Homer1-labeled postsynaptic puncta (green) in the CA1 *stratum radiatum* (left and middle). The merged image and expanded view on the right show excitatory synapses as co-localized puncta (yellow, white arrowheads). Scale bars, 20 µm and 2 µm (for expanded view) K, L. Quantification of co-localized puncta (excitatory synapses) per 100 µm^2^ (K) and their area covered (L). Data information: Data indicate mean ± SEM. Numbers on bars show tested cells or number of slices (G, L) and (number of animals). P-values are from unpaired Student’s t (H, I) or Mann-Whitney tests (D, E, K, L).

We first analyzed the density and morphology of CA1 apical dendritic spines, assuming we would find changes in the number of glutamatergic contacts (Fig 1B). Surprisingly, three-dimensional biocytin-based anatomical reconstructions did not reveal any changes in spine density or spine length in *Csf1r*^ΔFIRE/ΔFIRE^ mice compared to WT littermates (Figs 1C-E). In addition, morphometric characterization of CA1 pyramidal cells by Sholl analysis showed no changes in their dimension and dendritic arborization in *Csf1r*^ΔFIRE/ΔFIRE^ mice compared to WT littermates (Figs 1F-I). To extend our analysis to the whole synapse, we immunohistochemically labeled pre- and postsynaptic puncta with VGluT1 and Homer1, respectively. Consistent with the spine data, the number and size of excitatory synapses, as defined by the colocalization of both markers, were unchanged in the CA1 region in *Csf1r*^ΔFIRE/ΔFIRE^ compared with WT mice (Figs 1J-L). In summary, these findings indicate that excitatory synapses onto CA1 pyramidal cells are not reduced in number by the absence of microglia.

### Absence of microglia results in reduced excitability of CA1 pyramidal cells

We next investigated whether the absence of microglia has any consequences on the physiological function of CA1 neurons. Whole-cell patch-clamp recordings revealed no changes in input resistance, cell capacitance or in resting potential (Figs 2A-D). However, CA1 pyramidal cells produced a lower rate of action potentials on injection of depolarizing current in *Csf1r*^ΔFIRE/ΔFIRE^ mice compared to WT littermates (Figs 2E and F), along with an increase in minimal current to reach action potential threshold (rheobase) (Fig. 2G). No changes were seen in the threshold voltage eliciting action potentials (Fig 2H). This indicates lower neuronal excitability of CA1 pyramidal cells in *Csf1r*^ΔFIRE/ΔFIRE^ mice, which plausibly reflects changes in axonal electrophysiology, contrasting the lack of changes in passive membrane parameters, mainly defined by properties of the somatodendritic domain (Figs. 2B-D).

**Figure 2.**
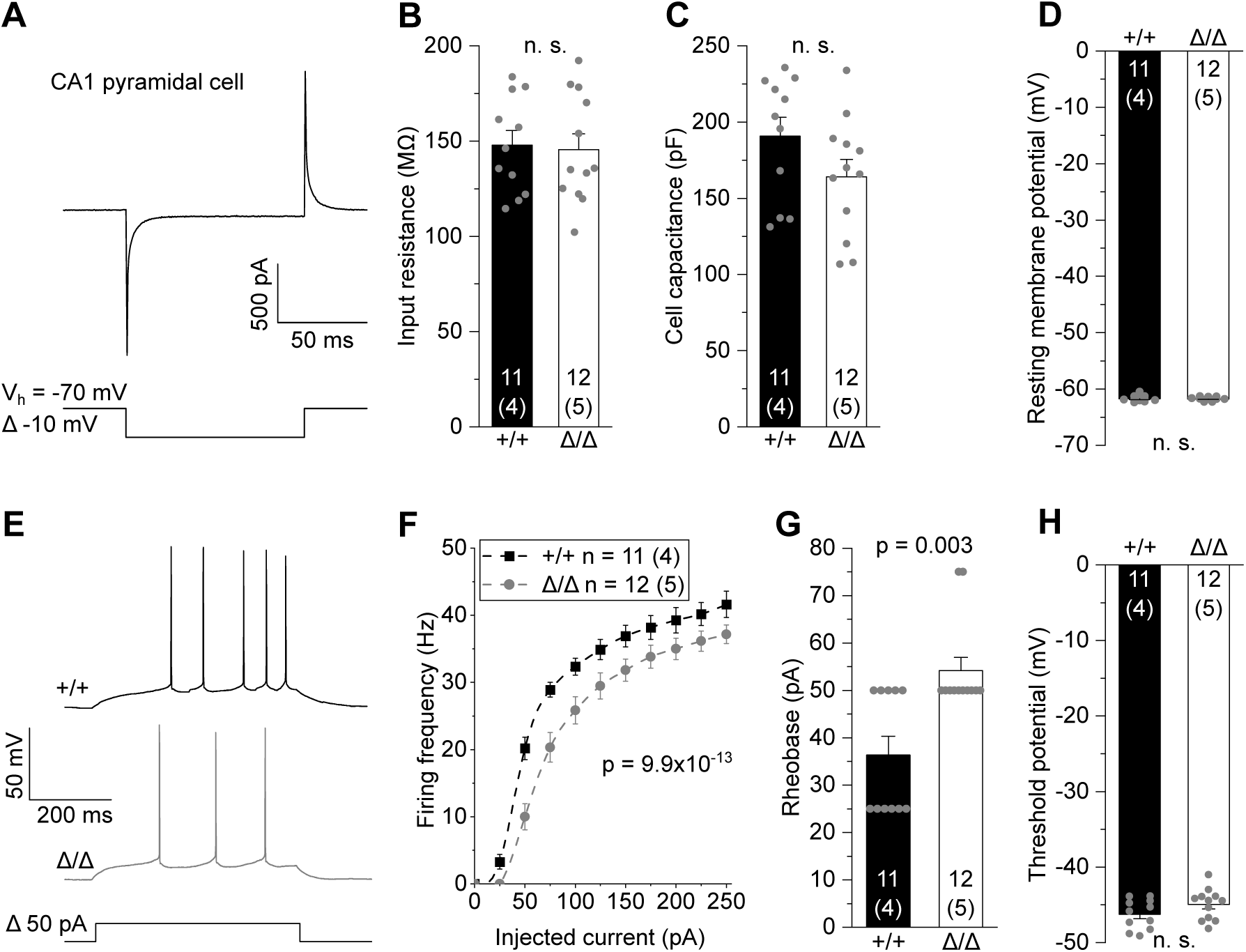
Absence of microglia results in reduced excitability of CA1 pyramidal cells. A. Patch-clamped membrane current of a CA1 pyramidal cell to a brief 10 mV hyperpolarization from which values for input resistance and cell capacitance were determined (see Materials and Methods). B-D. Quantification of membrane resistance (B), cell capacitance (C) and resting membrane potential (D) of CA1 pyramidal cells in Δ/Δ and +/+ mice. E. Specimen traces showing action potential firing patterns of CA1 pyramidal cells in response to 500 ms depolarizing current injections. F. Corresponding course of action potential firing frequencies on increasing depolarizations. G, H. Values for rheobase, i.e. minimal current to reach action potential threshold, (G) and action potential threshold voltage (H). Data information: Data are represented as mean ± SEM. Numbers on bars indicate tested cells and (number of animals). P-values are from unpaired Student’s t (B-D, H), or Mann-Whitney tests (G) and two-way ANOVA (F).

Notably, a similar picture was seen in heterozygous *Csf1r*^+/ΔFIRE^ mice, in which microglia were present at normal densities but were morphologically more ramified compared to WT littermates (Figs EV1A-C). As in microglial-deficient brains, no changes were seen in input resistance, cell capacitance and resting potential of CA1 pyramidal cells from *Csf1r*^+/ΔFIRE^ mice (Figs EV1D-F). Likewise, they were also less excitable compared to WT littermates, although the effect size was smaller than in homozygous *Csf1r*^ΔFIRE/ΔFIRE^ mice (Figs EV1G-J).

Taken together, these results demonstrate that the absence of microglia in *Csf1r*^ΔFIRE/ΔFIRE^ mice leads to a reduced ability of CA1 pyramidal cells to generate action potentials, thus limiting glutamatergic transmission downstream. These differences appear to be associated with microglial CSF1R signaling, as suggested by the qualitatively similar findings in heterozygous *Csf1r*^+/ΔFIRE^ mice with normal microglial density.

### Reduced CA3 – CA1 glutamatergic transmission in Csf1r^ΔFIRE/ΔFIRE^ mice

Based on the observed changes in neuronal output, we investigated the impact of microglial deficiency on hippocampal transmission focusing on excitatory inputs onto CA1 pyramidal cells. To this end, we electrically stimulated CA3 Schaffer collaterals at increasing stimulus strength and recorded compound excitatory postsynaptic currents (EPSCs) reflecting postsynaptic AMPA receptor (AMPAR) activation in CA1 pyramidal cells (Figs 3A and B). As revealed by the input-output relationships, *Csf1r*^ΔFIRE/ΔFIRE^ show greatly reduced EPSC amplitudes across the entire stimulation range compared to WT littermates (Fig 3C), indicating functional deficits in fast excitatory hippocampal transmission. We also observed reduced EPSC amplitudes in CA1 neurons in heterozygous *Csf1r*^+/ΔFIRE^ mice (Figs EV2A and B). Using the same recording configuration, we analyzed the amplitude ratio of two consecutive EPSCs in CA1 pyramidal cells by paired pulse stimulation of CA3 Schaffer collaterals. However, this revealed no changes in paired pulse ratio (PPR) in *Csf1r*^ΔFIRE/ΔFIRE^ (Figs 3D and E) as well as in *Csf1r*^+/ΔFIRE^ (Figs EV2C and D) compared to WT littermates.

**Figure 3.**
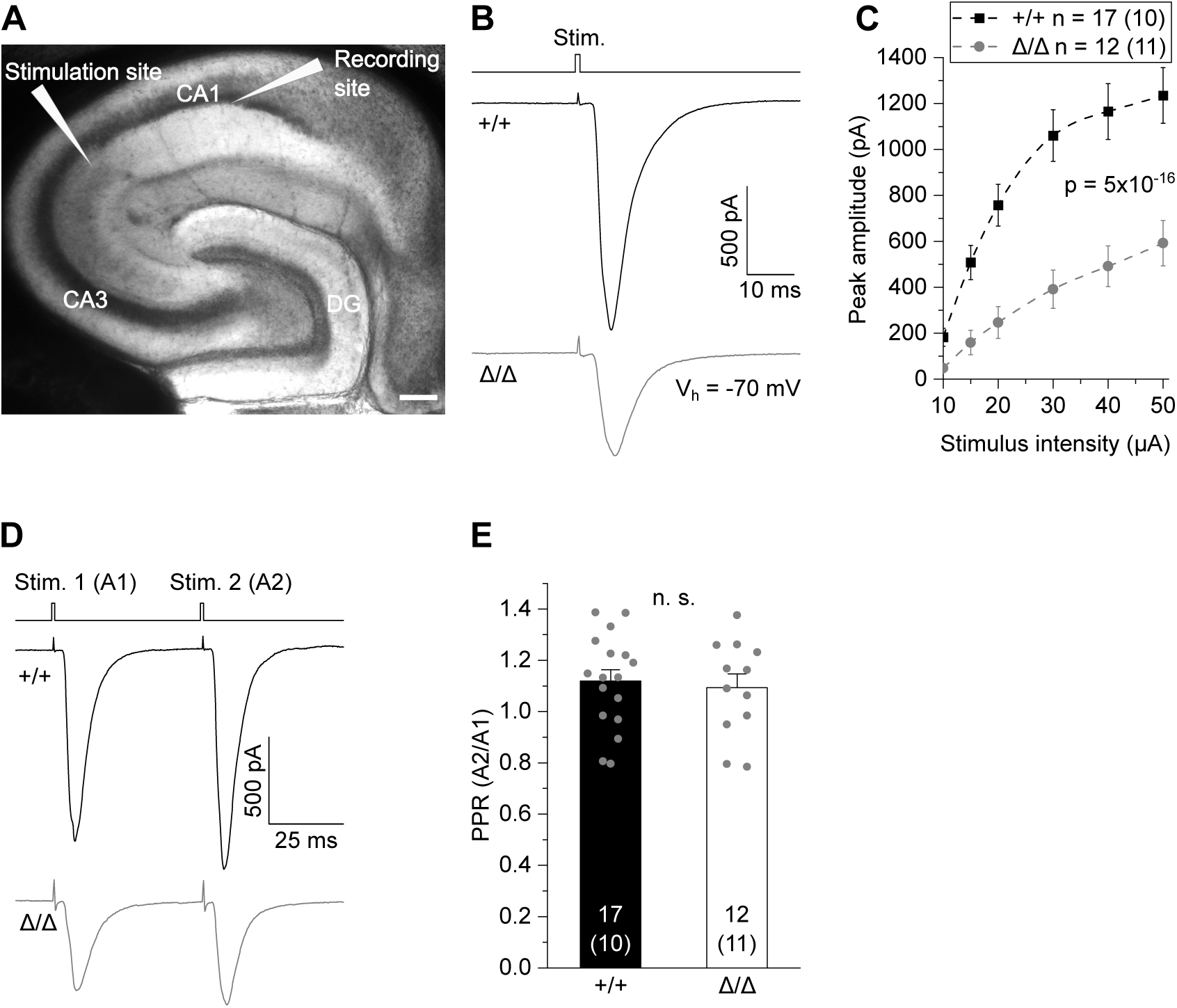
Reduced CA3 – CA1 glutamatergic transmission in *Csf1r*^ΔFIRE/ΔFIRE^ mice. A. Differential interference contrast image showing the localization of CA3 stimulation and CA1 recording sites. Scale bar, 200 µm. (CA1, CA3: *cornu ammonis regions 1 and 3*; DG: *dentate gyrus*) B. Specimen traces of excitatory postsynaptic currents (EPSC) in CA1 pyramidal cells in response to electrical stimulation for 0.2 ms. C. Corresponding input-output relationship showing peak amplitudes of CA1 EPSCs with increasing stimulation strength. D. Example traces (EPSC) in CA1 pyramidal cells after paired-pulse stimulation at 50 ms inter-stimulus intervals. E. Comparison of paired pulse ratios (PPR) as the quotient of the second vs first EPSC amplitude (A2/A1) of CA1 pyramidal cells. Data information: Data show mean ± SEM. Numbers on bars indicate tested cells and (number of animals). P-values are from two-way ANOVA (C) and unpaired Student’s t test (E).

Taken together, these results show that, although presynaptic glutamate release is preserved, the absence of microglia throughout development results in a lower excitatory input into CA1 pyramidal cells from the hippocampal CA3 region.

### Absence of microglia causes deficits in excitatory synaptic transmission

To test for changes in synaptic properties, we next analyzed spontaneous CA1 EPSCs (sEPSCs) including both action potential-dependent and -independent synaptic events, as well as action potential-independent miniature EPSCs (mEPSCs) in the presence of 0.3 µM tetrodotoxin (TTX) to block voltage-gated sodium channels (Fig 4A). Inter-event intervals of mEPSCs and sEPSCs were not different in *Csf1r*^ΔFIRE/ΔFIRE^ mice compared to WT littermates (Figs 4B and C), suggesting an unchanged number of functional glutamatergic synapses in agreement with our morphological findings. No changes were further observed in mEPSC decay time reflecting postsynaptic AMPAR-mediated currents (Figs 4D and E). Notably, analysis of current amplitudes showed impairment of multivesicular release, also referred to as synaptic multiplicity (i.e. the number of individual synaptic contacts and/or release sites between two connected neurons) in *Csf1r*^ΔFIRE/ΔFIRE^ mice as revealed by comparable peak amplitudes of sEPSCs and mEPSCs (Fig 4F). Given an unchanged number of functional synaptic contacts, this suggests a shift towards single-synaptic boutons with one presynapse contacting a single postsynaptic spine. In contrast, WT littermates exhibited larger amplitudes of sEPSC compared to mEPSC (Fig 4F), suggesting multiple presynaptic glutamate release sites per synapse as a characteristic feature of mature excitatory circuitry (Hsia et al., 1998).

**Figure 4.**
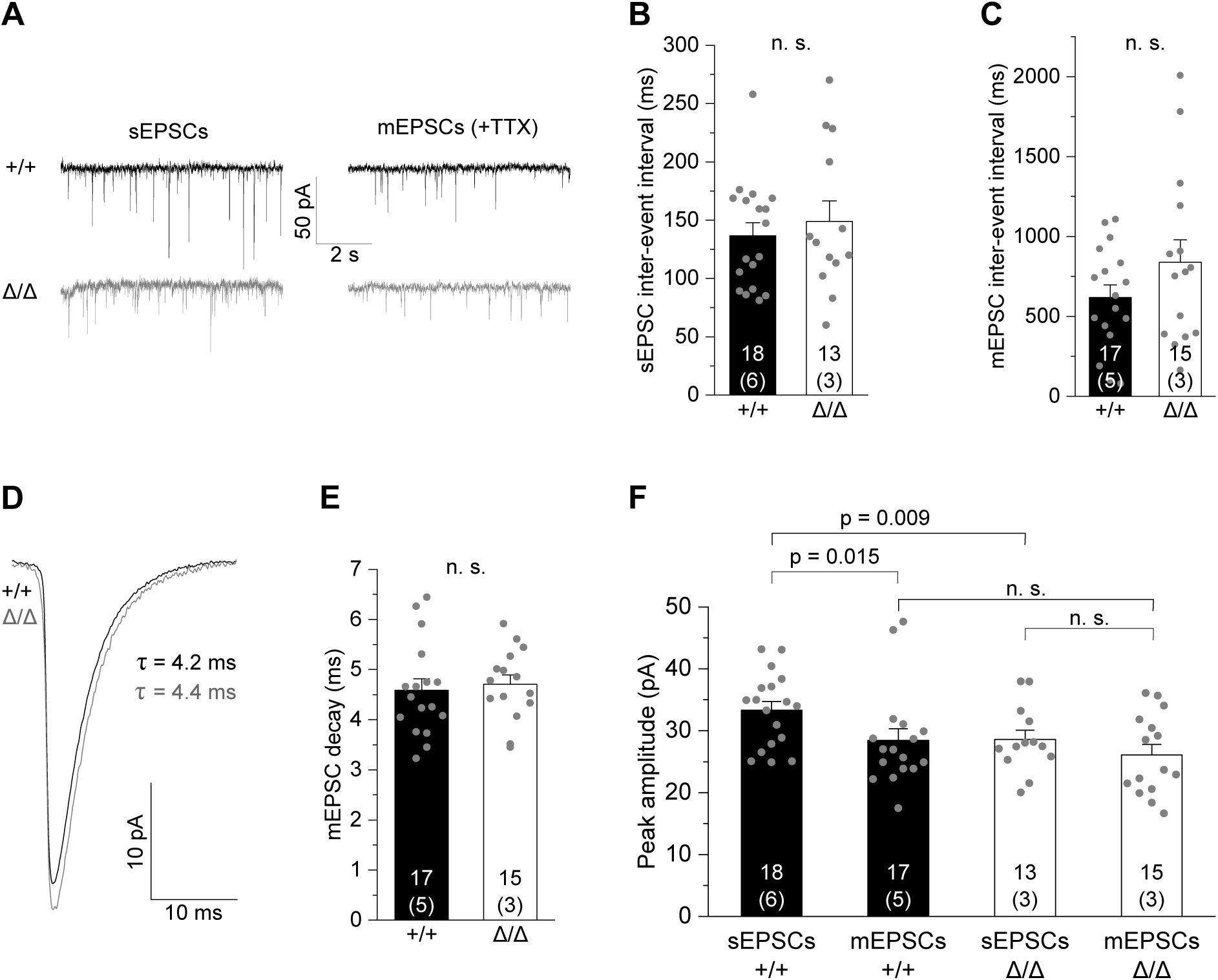
Absence of microglia causes deficits in excitatory synaptic transmission. A. Specimen traces showing AMPAR-mediated spontaneous EPSCs (sEPSCs) and miniature EPSCs (mEPSCs) in the presence of 300 nM TTX of CA1 pyramidal cells. B, C. Comparison of inter-event intervals of sEPSCs (B) and mEPSCs (C). D. Specimen traces showing average mEPSCs kinetics. E. Analysis of decay times of CA1 mEPSCs (AMPA receptor-evoked currents). F. Comparison of peak amplitudes of sEPSCs and mEPSCs. Note that mEPSC amplitudes are not different amongst genotypes, suggesting no change in functional synaptic contacts, whereas action potential-dependent sEPSC amplitudes are larger in +/+ but not in Δ/Δ mice, indicating impairment in synaptic multiplicity. Data information: Data indicate mean ± SEM. Numbers on bars show tested cells and (number of animals). P-values are from unpaired Student’s t [C, E, F (for Δ/Δ and sEPSC comparison)] or Mann-Whitney tests [B, F (for +/+ and mEPSC comparison)].

No changes were observed in GABAergic synaptic transmission since inter-event intervals and amplitudes of either spontaneous inhibitory postsynaptic currents (sIPSCs) or miniature inhibitory postsynaptic currents (mIPSCs) were unaltered in CA1 neurons of *Csf1r*^ΔFIRE/ΔFIRE^ mice (Figs EV3A-F).

Taken together, our findings suggest that microglia are dispensable for the pruning of excess synapses in hippocampal pyramidal neurons in *Csf1r*^ΔFIRE/ΔFIRE^ mice during brain development. However, the absence of microglia affects the maturation of glutamatergic synapses by impairment of multivesicular release.

### Absence of microglia leads to reduced synaptic NMDA receptor component in CA1 pyramidal cells

So far, the above experiments captured changes in fast excitatory transmission via activation of AMPARs (AMPAR), while NMDA receptor activation (NMDAR) was largely abolished due to its blockade by Mg^2+^ at negative membrane voltages. To investigate in more detail the changes in postsynaptic glutamate receptor function of both AMPAR and NMDAR, we adapted our recording conditions by perfusing slices with Mg^2+^-free extracellular solution in the presence of glycine and TTX, to promote NMDAR activation and shut down network activity. This resulted in dual-component mEPSCs carried by AMPAR plus NMDAR, and NMDAR-only mEPSCs (Fig 5A). To analyze the fraction of AMPAR vs NMDAR currents, expressed as electrical charge, we averaged >100 single events to create an average mEPSCs per cell and pharmacologically isolated the (i) AMPAR component from (ii) mixed AMPAR and NMDAR events by applying the specific NMDAR blocker D-AP5. Subtracting (ii) from (i) yields the calculated NMDAR component (Fig 5B). This revealed a lower NMDAR-mediated charge in *Csf1r*^ΔFIRE/ΔFIRE^ mice compared to WT littermates (Fig 5C) while, consistent with our findings above, no changes were seen for the AMPAR component (Fig 5D, cf. Figs 4E and F), resulting in a higher AMPAR/NMDAR charge ratio (Fig 5E).

**Figure 5.**
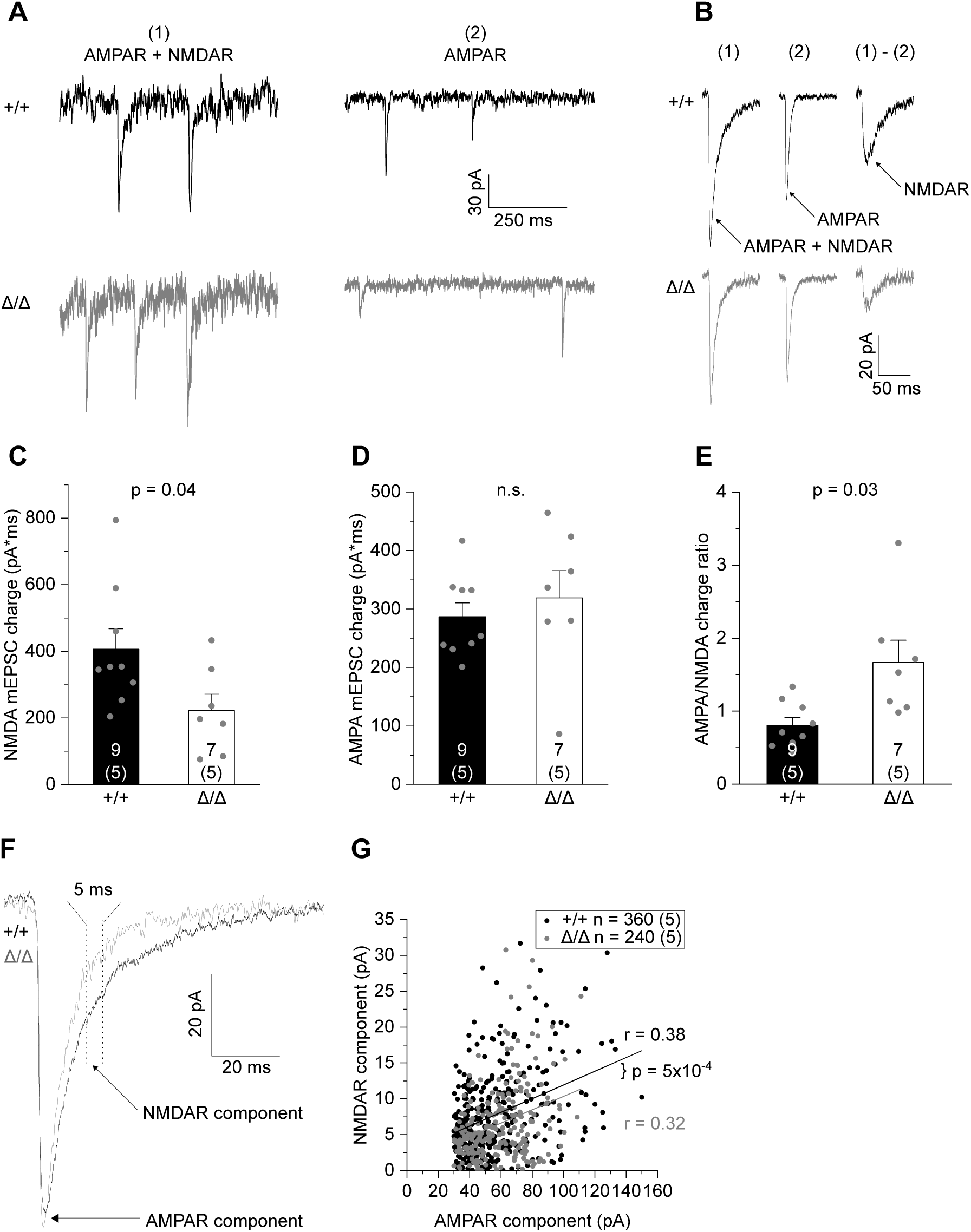
Absence of microglia results in reduced synaptic NMDA receptor component in CA1 pyramidal cells. A. Specimen traces showing dual-component mEPSCs comprising AMPAR- and NMDAR- evoked currents measured in nominally Mg^2+^-free extracellular solution in the presence of 300 nM TTX, 10 μM gabazine and 10 μM glycine (left), and pharmacologically isolated AMPAR-only mEPSCs after blockade of NMDARs with 50 µM D-AP5 (right). B. Average dual-component (1) and AMPAR-only (2) mEPSCs from which the synaptic NMDAR component was calculated by subtracting (1) – (2). C-E. Comparison of the NMDAR-(C) and AMPAR-mediated mEPSC charge (D) and the resulting AMPA/NMDA charge ratio (E). F. Specimen traces showing individual dual-component mEPSCs containing both AMPAR and NMDAR components. Arrows indicate where respective currents were measured. G. Correlation between the AMPA and NMDA measurements of individual mEPSCs comprising 40 events per cell. Data information: Data are represented as mean ± SEM. Numbers on bars indicate tested cells and (number of animals). P-values are from unpaired Student’s t test (C -E,) or ANCOVA (G, co-variant: AMPA peak amplitude). R-values refer to Pearson correlation coefficients.

Analysis of average mEPSCs assumes a homogeneous expression of glutamate receptors at individual postsynaptic sites, but there may be differences in the proportion of synapses in *Csf1r*^ΔFIRE/ΔFIRE^ mice with increased or decreased AMPAR and NMDAR function that may have been lost in averaging. To this end, we compared the AMPAR-mediated peak current of individual mEPSCs with its NMDAR current component (see Methods for details). Fitting of the data revealed a comparable degree of correlation of AMPAR to NMDAR currents in WT and *Csf1r*^ΔFIRE/ΔFIRE^ mice (Figs 5F and G). This suggests a similar distribution of synapses with different fractions of AMPAR versus NMDAR expression. In addition, analysis of covariance revealed a downward shifted regression line in *Csf1r*^ΔFIRE/ΔFIRE^ mice, corroborating our finding of a reduced NMDAR component at the level of individual synapses.

Due to the observed differences in synaptic NMDAR function in *Csf1r*^ΔFIRE/ΔFIRE^ mice, we extended our analysis to investigate tonic activation of NMDAR in CA1 pyramidal cells activated by ambient glutamate in the extracellular space. Tonic D-AP5-sensitive inward currents were measured in Mg^2+^-free, TTX-containing extracellular solution with glycine added as NMDAR co-agonist to unblock NMDAR and facilitate their activation (Fig EV4A). Under these conditions, all existing NMDAR-mediated currents contribute to the tonic current, comprising receptors located at mixed AMPAR/NMDAR synapses, silent synapses and extrasynaptically. Analysis of the D-AP5 sensitive current and RMS noise revealed an increase in both parameters in *Csf1r*^ΔFIRE/ΔFIRE^ mice compared to WT littermates (Figs EV4B and C), further indicating altered NMDAR function in mice deficient in microglia.

In summary, these data demonstrate the developmental impact of microglia in regulating postsynaptic function by a reduction of the NMDAR component.

### Absence of microglia changes cytokine and ApoE levels in the brain and results in reactive astrocytes

As the main immune cell type in the brain, microglia can affect neuronal function by releasing immune-modulatory signaling molecules, collectively referred to as cytokines. Notably, microglial cytokines play essential roles in synaptogenesis and synaptic transmission (Werneburg et al., 2017). To examine the consequences of permanent absence of microglia on cytokine levels in the forebrain (excluding olfactory bulb and cerebellum), we determined the levels of anti-and pro-inflammatory cytokines in *Csf1r*^ΔFIRE/ΔFIRE^ mice and WT littermates by high-sensitivity multiplex-ELISA in soluble and membrane-bound brain fractions (Fig 6A). *Csf1r*^ΔFIRE/ΔFIRE^ mice showed subtle changes in cytokine levels with increases in anti-inflammatory IL-10 and IL-4, while pro-inflammatory IFNγ and IL-2 were reduced (Fig 6B and Table 1). Notably, TNFα and IL-1ß levels, two key pro-inflammatory microglial cytokines, were unchanged in the brains of *Csf1r*^ΔFIRE/ΔFIRE^ mice compared to *Csf1r*^+/ΔFIRE^ and WT littermates (Fig 6B and Table 1). In addition, *Csf1r*^+/ΔFIRE^ mice revealed subtle reductions in INFγ as well as CXCL1 (Table 1).

**Figure 6.**
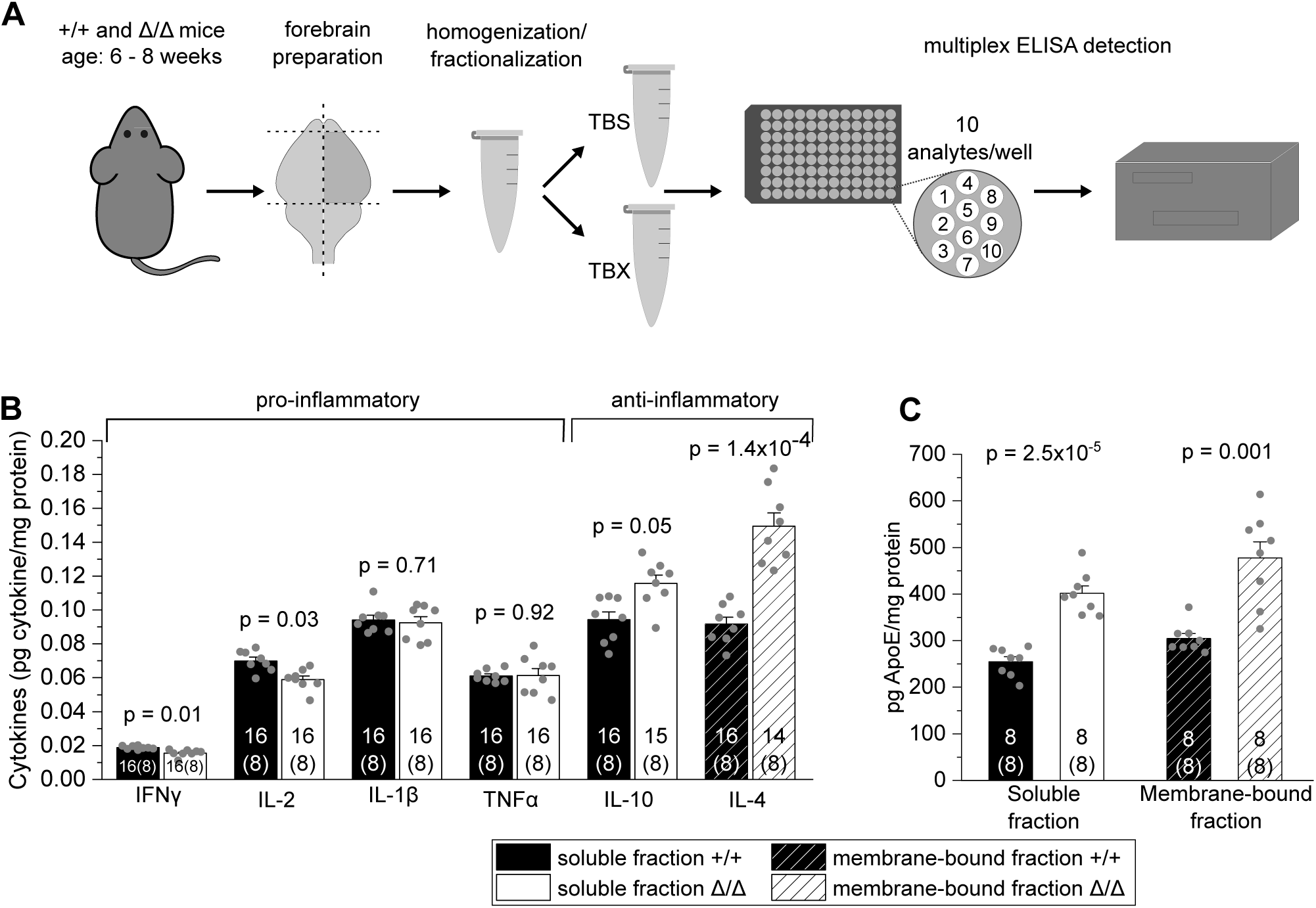
Absence of microglia changes cytokine and ApoE levels in the brain. A. Workflow illustrating the ELISA-based detection of brain cytokines and ApoE in soluble (TBS) and membrane-bound (TBX) fractions from cortical brain homogenates (see Materials and Methods). B, C. Comparison of basal levels of anti- and pro-inflammatory cytokines (B) and apolipoprotein E (ApoE) (C) in soluble and membrane-bound fractions in Δ/Δ mice and +/+ mice. Data information: Data are represented as mean ± SEM. Numbers on bars indicate technical replicates and (number of animals). P-values are from unpaired Student’s t tests corrected for multiple comparisons.

**Table 1.**
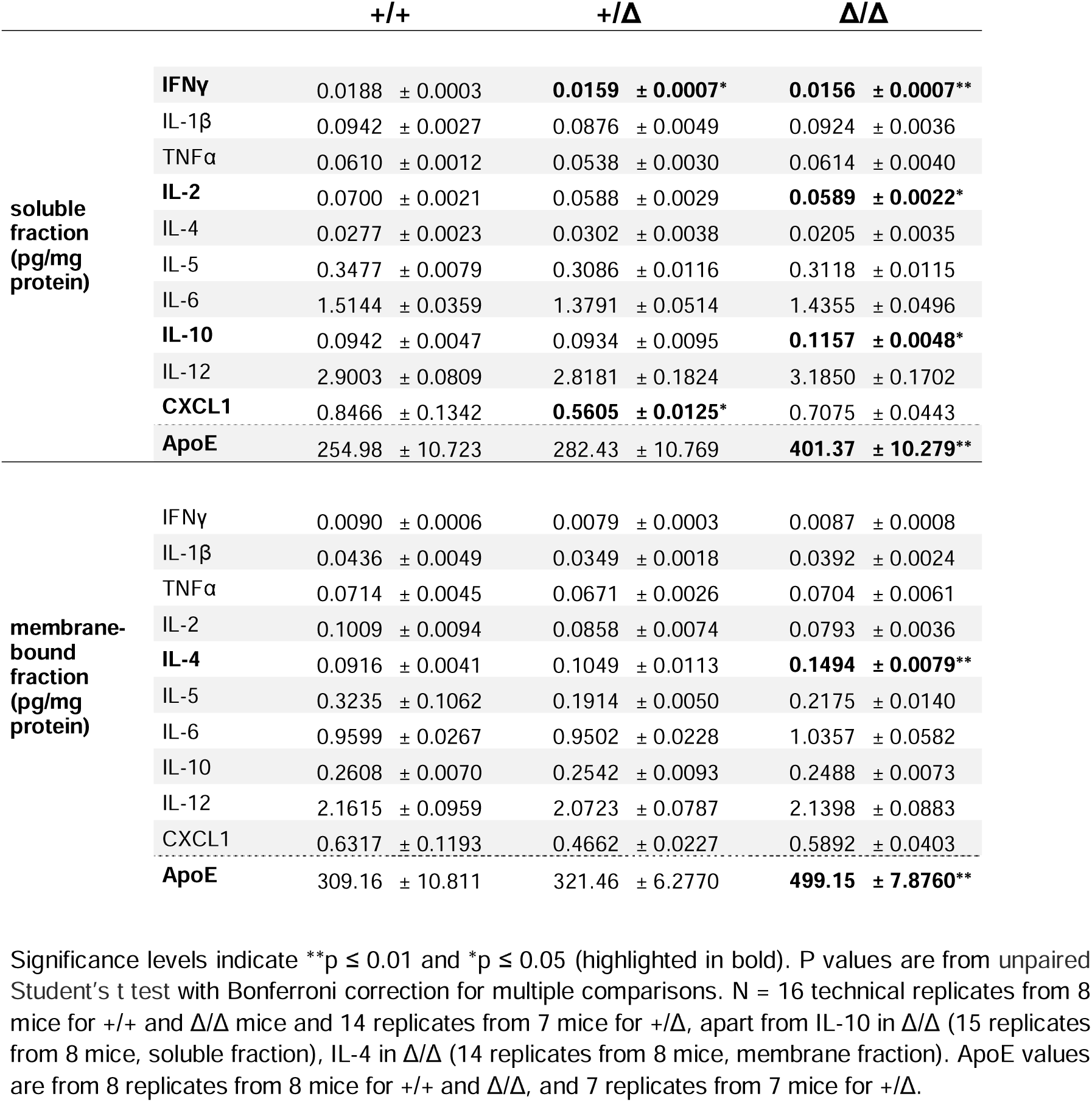
Basal cytokine levels in the brains of *Csf1r^+/+^*, *Csf1r^+/^*^Δ^ and *Csf1r*^Δ*/*Δ^ mice.

Apolipoprotein E (ApoE) is mainly produced by astrocytes in the murine brain (Boyles et al., 1985; Pitas et al., 1987; Zhang et al., 2014 and 2016). Interestingly, soluble and membrane bound ApoE levels were strongly increased in the brains of *Csf1r*^ΔFIRE/ΔFIRE^ mice compared to *Csf1r*^+/ΔFIRE^ and WT littermates (Fig 6C and Table 1), suggesting that changes in ApoE result from the absence of microglia rather than alterations in CSF1R function. Morphological analysis of astrocytes in the CA1 stratum radiatum of *Csf1r*^ΔFIRE/ΔFIRE^ mice revealed an increase in both intensity and area covered by GFAP immunoreactivity, while astrocyte numbers remained unchanged compared to WT littermates (Figs 7A-D). These changes are also reflected in increased complexity of astrocyte morphology, as evidenced by Sholl analysis revealing increases in the number of intersections and total process length (Figs 7E-H).

**Figure 7.**
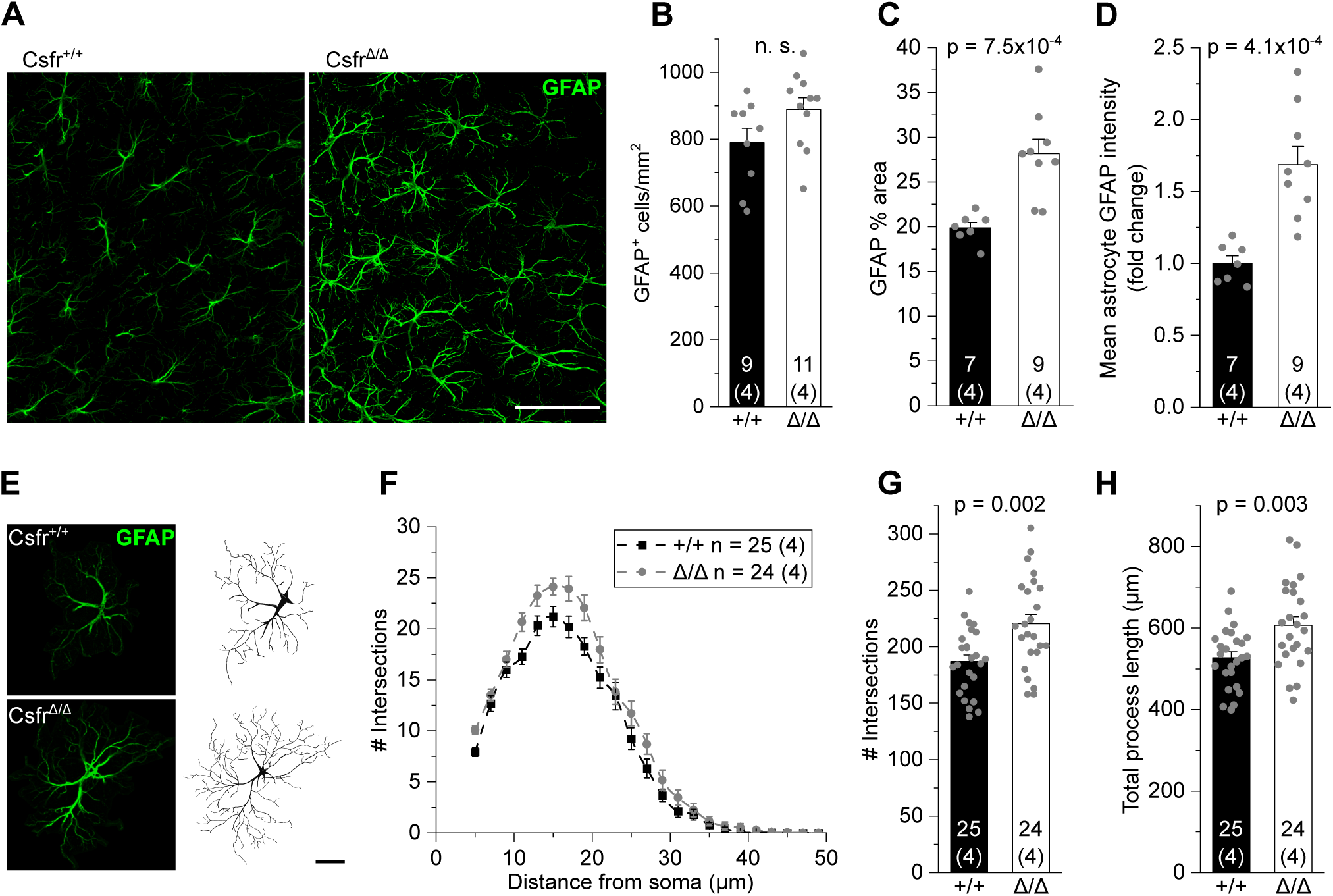
Absence of microglia results in reactive astrocytes. A. Specimen confocal images illustrating astrocytes by GFAP immunoreactivity (green) in the CA1 *stratum radiatum*. Scale bar, 50 µm. B-D. Analyses of GFAP^+^ cell density (B), percentage of area covered by GFAP^+^ astrocytes (C) and mean GFAP intensity (D). E. Specimen confocal images illustrating astrocyte morphology by GFAP immunoreactivity (left) and their 3D-reconstruction (right). Scale bar, 10µm. F-H. Sholl analysis-derived values of the number of intersections with Sholl radii at increasing distance from the soma (F) and resulting total number of intersections (G) and total process length (H) of astrocytes. Data information: Data are represented as mean ± SEM. Numbers on bars indicate number of slices and (number of animals). P-values are from unpaired Student’s t tests (B-D, G, H).

Taken together, these data indicate a trend toward an anti-inflammatory state in brains of *Csf1r*^ΔFIRE/ΔFIRE^ mice. This is accompanied by reactive astrocyte morphology and elevated ApoE levels, suggesting functional changes in astrocytes in the absence of microglia in *Csf1r*^ΔFIRE/ΔFIRE^ mice.

### Absence of microglia does not alter object recognition memory

Finally, we asked whether the observed functional changes are reflected at the behavioral level by affecting memory. Novel object recognition is a widely accepted task for assessing nonspatial memory in rodents with an important contribution of the hippocampus and perirhinal cortex (Cohen and Stackman, 2015). Here, after habituation to a familiar object, mice are exposed to a novel object to interact with. We detected no differences in the performance in the novel object recognition test of 6-10-week-old male and female *Csf1r*^ΔFIRE/ΔFIRE^ mice compared to *Csf1r*^+/ΔFIRE^ and WT littermates (Figs 8A and B), thus excluding nonspatial memory impairments in the absence of microglia.

**Figure 8.**
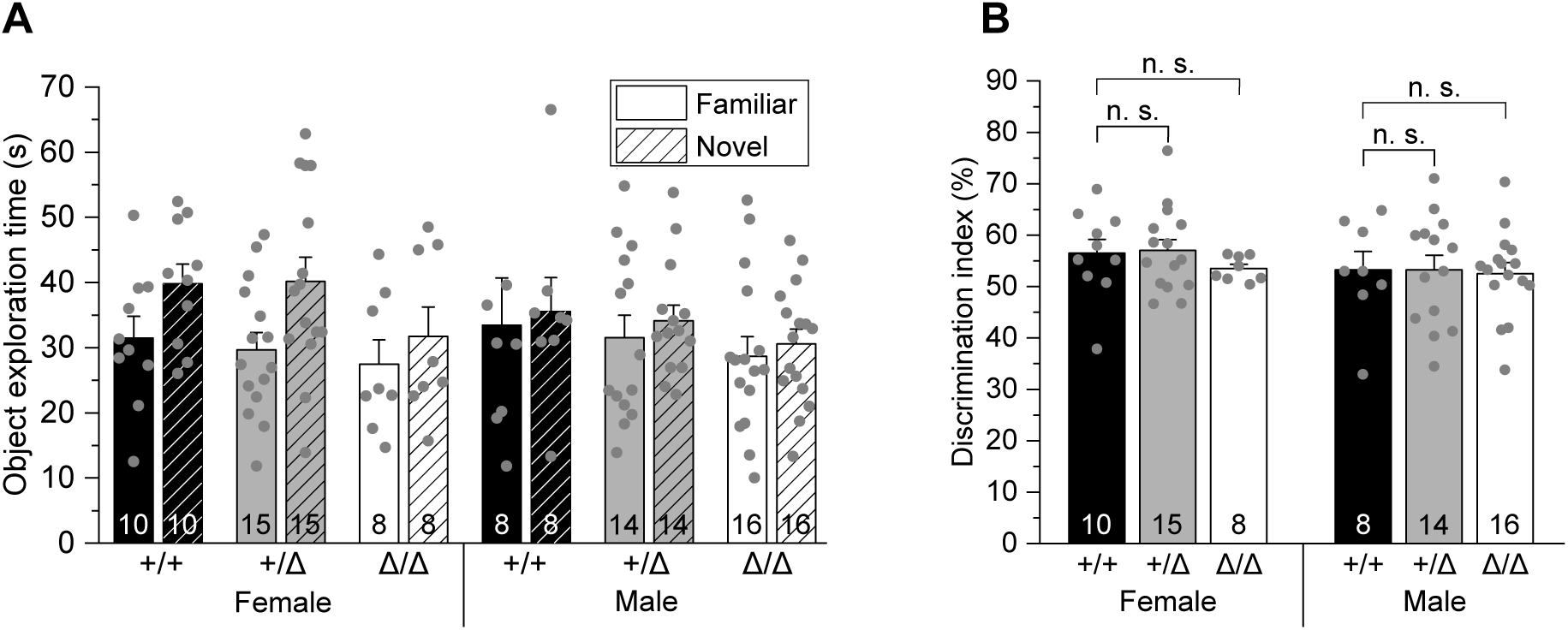
Absence of microglia does not alter object recognition memory. A. Comparison of exploration times of the familiar and novel object after habituation for each genotype and sex (see Materials and Methods). B. Comparison of the respective discrimination indices, defined as the time exploring the novel object / time exploring the novel and familiar object *100. Data information: Data are represented as mean ± SEM. Numbers on bars indicate number of tested animals. P-values are from two-way ANOVA (B)

## Discussion

In this study, we investigated the role of microglia in embryonic and postnatal development with regard to their capacity to sculpt glutamatergic circuitry in the hippocampus and contribute to nonspatial memory. We examined synapse density and dendritic spine morphology along with the electrophysiological profile of hippocampal pyramidal neurons on the synaptic and cellular level, and cognitive function in the NOR task in young adult *Csf1r*^ΔFIRE/ΔFIRE^ mice that are deficient in microglia throughout life. Our motivation was further fueled by the fact that these mice develop surprisingly normal without abnormalities in behavioral tasks (Rojo et al., 2019; McNamara et al., 2023), which appears to contradict findings attributing a crucial role to microglia in neural development and function.

Our study has disclosed the following main results: (1) Number and size of excitatory synapses in the CA1 *stratum radiatum*, as well as spine density of apical dendrites of CA1 pyramidal cells are unchanged in *Csf1r*^ΔFIRE/ΔFIRE^ mice, suggesting that microglia are dispensable for synaptic pruning. (2) CA1 pyramidal cells are less excitable and receive less excitatory input from CA3 Schaffer collaterals in *Csf1r*^ΔFIRE/ΔFIRE^ mice, resulting in a weakened glutamatergic transmission in the absence of microglia. At the synaptic level, this is accompanied by impairment of multivesicular release and a reduction in the postsynaptic NMDA receptor component, without changes in inhibitory GABAergic transmission. (3) Nonspatial memory, to which hippocampus and perirhinal cortex contribute, is not impaired in *Csf1r*^ΔFIRE/ΔFIRE^ mice. (4) Absence of microglia results in a cerebral environment with mild increases in anti-inflammatory cytokines along with strong increases in ApoE levels and GFAP area and intensity, indicating astrocyte reactivity.

### Dispensability of microglia to eliminate the surplus of synapses of developing hippocampal neurons

A major outcome of our work is that the developmental and postnatal absence of microglia in young adult *Csf1r*^ΔFIRE/ΔFIRE^ mice does not affect synaptic markers and spine density in apical dendritic regions of CA1 pyramidal cells. This is a surprising finding given that microglia have been suggested to prune the majority of excess synapses during the first postnatal weeks in order to form a properly wired and adaptive brain (Schafer and Stevens, 2015). Spine dynamics of hippocampal CA1 neurons remain extremely high also beyond this pruning period with a turnover of spines every 1-2 weeks in 10-12-week-old mice (Attardo et al., 2015; Pfeiffer et al., 2018), a time still captured in our analysis. This implies that even minor microglial influences on CA1 synapse elimination or remodeling would have been expected to produce a visible effect in the number of spines in *Csf1r*^ΔFIRE/ΔFIRE^ mice. Nevertheless, we observed no morphological changes in synapse area and spine morphology, suggesting normal physical interactions between pre- and postsynaptic sites.

None of the previous studies providing evidence for microglial-dependent synapse elimination during postnatal development in hippocampus, thalamus and visual cortex (Paolicelli et al, 2011; Schafer et al., 2012; Zhan et al., 2014; Sipe et al., 2016; Filipello et al., 2018; Liu et al. 2021) investigated the consequences of permanent absence of microglia during embryonic development and postnatal life on neuronal development and function. Our findings in *Csf1r*^ΔFIRE/ΔFIRE^ mice, which in addition do not involve potential confounding factors such as cell ablation, suggest that microglia are dispensable for the regulation of the number of synapses post-developmentally, consistent with recent findings showing that microglia in the developing hippocampus do not phagocytose postsynaptic material (Weinhard et al., 2018).

Microglial pruning is not a random process but steered by synaptic activity (Tremblay et al., 2010, Schafer et al., 2012, Gunner et al., 2019), leading to the preferential elimination of weaker connections while sparing the stronger ones. Indeed, we find weaker and more immature synaptic connections in the hippocampal glutamatergic network in *Csf1r*^ΔFIRE/ΔFIRE^ mice. However, this is unlikely to result from a deficit in elimination of weaker synaptic contacts, which would have caused an increase in spine density. Instead, the absence of microglia impairs the maturation of glutamatergic synapses (further discussed below). These synaptic changes seem to persist into adulthood, as observed in adult mice with genetically deleted microglial CR3, CX3CR1 and TREM2 signaling (Paolicelli et al., 2011, Zhan et al., 2014, Filipello et al., 2018).

### Role of microglia-independent mechanisms in synaptic pruning

Which alternative cell types or mechanisms may regulate the pruning of synapses in brains devoid of microglia? Around the same time as microglia, astrocytes were identified as critical regulators of the pruning of synapses in the developing and adult brain (Chung et al., 2013). Microglia also cooperate with astrocytes to control the number of synapses, for example via specific cytokine-mediated interactions (Lim et al., 2013, Vainchtein et al., 2018), and to clear apoptotic neurons through phagocytic uptake (Damisah et al., 2020). Moreover, microglia can eliminate newborn astrocytes via phagocytic removal, thereby indirectly affecting the impact of astrocytes on synapse regulation (VanRyzin et al., 2019). Indeed, our findings suggest astrocyte reactivity, as revealed by increases in GFAP area and intensity, as well as elevated levels of ApoE and IL-10 in *Csf1r*^ΔFIRE/ΔFIRE^ mice (Boyles et al., 1985; Zhang et al., 2014; Villacampa et al., 2015). While IL-10 plays a role in synapse formation (Lim et al., 2013), ApoE controls the rate of synaptic pruning by astrocytes (Chung et al., 2016) and can affect glutamatergic transmission in multiple ways via signaling through synaptic ApoE receptors (Lane-Donovan and Herz, 2017). It is tempting to speculate that astrocytes may at least partially compensate for the loss of microglia in *Csf1r*^ΔFIRE/ΔFIRE^ mice, consistent with reports showing an up-regulation of their phagocytic capacity in mice with dysfunctional or ablated microglia (Konishi et al., 2020; Berdowski et al., 2022).

Future studies will help to determine what proportion of excess or unwanted synapses in development are subject to glial elimination as opposed to other processes, whether and how these processes differ between brain regions, and which specific functional properties of synapses determine their fate. As previously suggested, the simplified view that only weaker synaptic contacts are removed falls short of reflecting the high degree of structural and functional heterogeneity of synapses in neural networks (Wichmann and Kuner, 2022).

### Microglia affect glutamatergic network maturation in the hippocampus

The absence of microglia during embryonic development and postnatal life resulted in a distinct electrophysiological phenotype in *Csf1r*^ΔFIRE/ΔFIRE^ mice, with weakened and immature glutamatergic hippocampal transmission compared to WT littermates. CA1 pyramidal cells in *Csf1r*^ΔFIRE/ΔFIRE^ mice received less excitatory input from the CA3 region and are less capable of generating excitatory output. While presynaptic glutamate release probability was preserved, we identified changes in pre- and postsynaptic function, as evidenced by impairment of multivesicular release and reduction of NMDAR-mediated postsynaptic charge transfer. In contrast, we found no changes in inhibitory GABAergic synaptic input into CA1 pyramidal neurons in Csf1r^ΔFIRE/ΔFIRE^ mice.

Interestingly, the electrophysiological profile in *Csf1r*^ΔFIRE/ΔFIRE^ mice closely mirrors the changes in hippocampal transmission that emerged after transient depletion of microglia in adult mice that had undergone normal development (cf. Table 2 for a detailed comparison). In these cases, only excitatory transmission was impaired, with CA1 pyramidal cells receiving less glutamatergic input from CA3 Schaffer collaterals, while inhibition also remained unaffected. Almost congruent with our findings in *Csf1r*^ΔFIRE/ΔFIRE^ mice, this was related to impairment in multivesicular release and changes in postsynaptic NMDA receptors without affecting the AMPAR component (Parkhurst et al., 2013; Basilico et al., 2022; Ma et al., 2020). The observed reduction in the postsynaptic NMDAR component may be related to an increase in ambient glutamate levels, as suggested by the increase in tonic D-AP5-sensitive inward current in *Csf1r*^ΔFIRE/ΔFIRE^ mice. In consequence, NMDAR desensitization would be increased, leaving fewer NMDARs available for synaptic activation (Cavelier et al., 2005). Notably, microglial depletion seems to affect glutamatergic and GABAergic transmission differentially in the visual cortex compared to hippocampus and motor cortex (Table 2).

**Table 2.** Similar electrophysiological profile in the hippocampus of *Csf1r*^ΔFIRE/ΔFIRE^ mice compared to data from wild type mice with microglial depletion at comparable ages.

|  | Present study | Basilico et al., 2022 | Du et al., 2022 | Parkhurst et al., 2013 | Liu et al., 2021 |
| --- | --- | --- | --- | --- | --- |
| brain region | hippocampus CA1/3 | hippocampus CA1/3 | hippocampus CA1/3 | motor cortex L5 | visual cortex L5 |
| age | 6-10 weeks | 6-10 weeks | 4-5 weeks | 3-6 weeks | 9-10 weeks |
| microglial depletion | throughout life, 100% | transient (≈ 85-90 %), postnatally/adolescence |  |  |  |
| **Glutamatergic transmission** | ↓ | ↓ | ↓ | ↓ | ↑ |
| • I/O (CA3 - CA1) | ↓ | ↓ | ↓ | n.d. | n.d. |
| • release probability | □ □ | ↑ | □ | n.d. | n.d. |
| • mEPSC (AMPA) | freq □ amp □ | freq □ amp □ | n.d. | freq ↓ amp □ | n.d. |
| • multivesicular release | ↓ | ↓ | n.d. | n.d. | n.d. |
| • NMDAR (postsyn.) | ↓ | ↓ | n.d. | ↓ | n.d. |
| **GABAergic transmission** | □ | □ | n.d. | n.d. | ↑ |
| • m/sIPSCs | □ | □ | n.d. | n.d. | n.d. |

Key to symbols and abbreviations:
↓ / ↑ / – , increase / decrease / no change; n.d., not determined.
mEPSC, miniature excitatory postsynaptic current; m/sIPSC, miniature/spontaneous inhibitory postsynaptic current; freq, frequency; amp, amplitude; I/O, Input/Output relationship referring to CA1 compound EPSCs in response to CA3 electrical stimulation; AMPA, alpha-amino-3-hydroxy-5-methyl-4-isoxazolpropionic acid; NMDAR, N-methyl-D-aspartate receptor; GABA, gamma-aminobutyric acid.

A notable feature in both *Csf1r*^ΔFIRE/ΔFIRE^ and microglia-depleted WT mice is the impairment of multivesicular release (Basilico et al., 2022, Table 2), a hallmark of mature hippocampal synaptic transmission (Hsia et al., 1998, Rigby et al., 2022). This may be due to a preferential interaction of microglia with presynaptic structures, contributing to the formation of multi-synaptic boutons in the developing hippocampus (Zhan et al., 2014, Weinhard et al., 2018). Related to this, activated microglia were shown to displace presynaptic terminals in a process called synaptic stripping (Trapp et al., 2007; Chen et al., 2014).

Apart from the effects in microglial-deficient mice, we also observed lower CA1 excitability and EPSC amplitude in heterozygous *Csf1r*^+/ΔFIRE^ mice, in which microglia are present without changes in cell density (Munro et al., 2020). Similarly, a reduced excitatory input into CA1 pyramidal cells was reported in mice with fully deleted fractalkine receptor signaling, a main pathway for microglia-neuron interactions (Basilico et al., 2019). It should be noted that both *Csf1r*^ΔFIRE/ΔFIRE^ and *Csf1r*^+/ΔFIRE^ mice are on a mixed genetic background which may affect the results and underscore the need to compare with lines from different breedings.

Despite the lack of overt behavioral abnormalities in *Csf1r*^ΔFIRE/ΔFIRE^ mice (Rojo et al., 2019), an obvious assumption related to the altered electrophysiological profile of CA1 pyramidal cells are alterations in hippocampal-related behavior. However, we observed no differences in nonspatial memory in *Csf1r*^ΔFIRE/ΔFIRE^ mice. Importantly, spatial learning and memory encoding were also normal in *Csf1r*^ΔFIRE/ΔFIRE^ mice (McNamara et al., 2023). These findings are reminiscent of a previous study depleting microglia in young wild type mice, which also resulted in no behavioral deficits, in fact even slight improvements, in the Barnes maze and contextual fear conditioning (Elmore et al., 2014).

Finally, our data suggest changes in astrocytes as a result of the absence of microglia in *Csf1r*^ΔFIRE/ΔFIRE^ mice. These include a reactive phenotype and a strong increase in ApoE levels in both soluble and membrane-bound brain fractions of *Csf1r*^ΔFIRE/ΔFIRE^ mice, suggesting increased release from astrocytes as the main producers of ApoE in the brain (Boyles et al., 1985; Pitas et al., 1987; Zhang et al., 2014 and 2016). Due to its paramount role in brain lipid metabolism and the ability to suppress the release of pro-inflammatory cytokines (Laskowitz et al., 1997 and 1998; Flowers and Rebeck 2020), the increase in ApoE is expected to promote anti-inflammatory processes and neural homeostasis. Indeed, we observed increases in IL-4 and IL-10 along with decreases in INF-γ and IL-2 in the brains of *Csf1r*^ΔFIRE/ΔFIRE^ mice. It is important to consider that the observed changes reflect the situation in the healthy brain, which is likely to be different in the presence of brain damage or disease, in which the permanent absence of microglia has been associated with accelerated pathology in a dementia model and in prion disease (Bradford et al., 2022; Shabestari et al., 2022).

Collectively, our findings provide evidence that microglia are dispensable for synaptic pruning in the developing hippocampus and suggest that microglial-independent mechanisms are capable of achieving a basic, albeit not fully mature, network connectivity without causing behavioral disturbances. However, microglia contribute to fine-tune the maturation and strength of glutamatergic synapses within the hippocampal network, a role they seem to maintain in adulthood.

## Materials and Methods

### Mice

Experiments used transgenic *Csf1r*^ΔFIRE/ΔFIRE^ mice (Rojo et al., 2019) which were on a mixed C57BL/6J x CBA/J background with littermate heterozygous *Csf1r*^ΔFIRE/+^ and littermate wild-type *Csf1r*^+/+^ controls of both sexes aged 6-10 weeks. Mice were housed in groups in individually ventilated cages under specific-pathogen-free conditions on a 12-h light/dark cycle with food and water *ad libitum.* All procedures involving handling of living animals were carried out in accordance with the UK and German animal protection law and approved by the local authorities for health and social services (LaGeSo T-CH 0043/20).

### Novel object recognition test

The novel object recognition (NOR) test was performed in a dedicated behavioral testing room using mice of both sexes, with females and males always being tested on different days and analyzed separately. This test was performed on mice at 1-2 months of age to assess nonspatial memory. A total of 24 *Csf1r*^Δ*FIRE/*Δ*FIRE*^ mice (16 male, 8 female), 29 *Csf1r*^Δ*FIRE/+*^ littermates (14 male, 15 female), and 18 *Csf1r^+/+^* littermates (8 male, 10 female) were tested. Mice were housed under standard conditions, with 12-hour light/dark cycles, in temperature and humidity-controlled rooms. Researchers were blinded to experimental groups and remained blinded during data analysis. Handling was carried out 3-4 days prior to testing to reduce handling-induced stress on testing days. Equipment was cleaned with 70% ethanol between tests. Before starting this test, mice were run on an open field (47 cm x 47 cm), which habituated them to the arena. Mice were then placed back in the open field on a separate day and habituated to two identical objects, Lego towers. The following day (the testing day), one Lego tower was replaced by a light bulb as a novel object and mice were left to freely explore the novel and familiar objects for 10 min. Analyses of the acquired behavior videos were performed using Any-maze software. Exploration of an object was timed when the head of the mouse was within 1 cm of the object. Exploration time was not counted when the mouse was grooming near the object (it was only counted when the mouse was sniffing, examining, touching, or climbing on the object). No outliers were removed.

### Brain slice preparation

Mice were decapitated under isoflurane anesthesia. Whole brains were rapidly removed from the skull and immediately immersed in ice-cold slicing solution. Acute horizontal brain slices (300 μm) of the ventral hippocampus were prepared according to the protocol of Bischofberger et al. (2006) using a sucrose-based slicing solution containing (mM): 87 NaCl, 25 NaHCO_3_, 2.5 KCl, 0.5 CaCl_2_, 3 MgCl_2_, 1.25 NaH_2_PO_4_, 10 glucose, 75 sucrose, pH 7.4, bubbled with 95%O_2_ / 5%CO_2_ at < 4°C. After cutting, slices were allowed to recover for 30 min in warmed (34-36°C) slicing solution and then kept at room temperature until experimental use.

### External and intracellular solutions

For all experiments, slices were superfused with bicarbonate-buffered artificial cerebrospinal fluid (ASCF) at 34-36°C containing (mM): 124 NaCl, 2.5 KCl, 26 NaHCO_3_, 1 NaH_2_PO_4_, 2 CaCl_2_, 1 MgCl_2_, 10 glucose, bubbled with 95%O_2_ / 5%CO_2_.

For excitatory currents, cells were recorded with a potassium gluconate-based intracellular solution containing (mM): 120 K-gluconate, 10 HEPES, 10 EGTA, 2 MgCl_2_, 5 Na_2_-phosphocreatine, 2 Na_2_ATP, 0.5 Na_2_GTP, 5 QX-314 Cl or KCl (for excitability measurements). In some experiments, 0.1% Biocytin was added for post-hoc anatomical visualization and reconstruction of neuronal morphology.

For inhibitory currents, a KCl-based intracellular solution was used containing (mM): 125 KCl, 4 NaCl, 10 HEPES, 10 EGTA, 1 CaCl_2_, 4 MgATP, 0.5 Na_2_GTP. Intracellular solutions were adjusted to a final osmolarity of 285±5 mOsmol/L and a pH of 7.2.

### Electrophysiology

Hippocampal neurons in slices were visualized by IR-DIC optics using an upright Scientifica SliceScope equipped with a 63x water immersion objective (N.A. 1.0) and Olympus XM10 camera.

Whole-cell recordings from pyramidal neurons were obtained at a depth of > 40 μm below the slice surface using borosilicate glass pipettes with a tip resistance of 2.5 - 3.5 MΩ, resulting in access (series) resistances of < 20 MΩ that were compensated by ∼60%. Cells with changes in series resistance > 20% were excluded from the analysis. Voltage- and current-clamp recordings were performed using a Multiclamp 700B amplifier (Molecular Devices). Currents were filtered at 2 kHz (10 kHz for membrane test), digitized (20 kHz) and analyzed off-line using pClamp10 software. Resting membrane potential was measured in current clamp mode immediately after breaking into the cell. Electrode junction potentials were not compensated and were 14.5 mV and 3 mV for K-gluconate- and KCl-based intracellular solutions, respectively (determined using the *LJP* tool in pClamp10).

Input (Rt) and series resistances (Rs) were analyzed from voltage clamped cells held at −70 mV by applying 10 mV hyperpolarizing voltage steps using Ohm’s law (R=V/I). Total (input) resistances (Rt) and (Rs) were calculated as follows: Rt = V (applied voltage step) / Is (steady-state current), Rs = V (applied voltage step) / Ip (transient peak capacitive current). Cell capacitance (Cm) was calculated from the time constant (tau) of the current transient decay using a mono exponential fit according to Cm = tau/(Rs*Rm/Rs+Rm) = tau*(Is+Ip)^2^/(V*Is).

For determining neuronal excitability, CA1 pyramidal cells were held in current clamp mode and voltage responses filtered and digitized at 10 and 20 kHz, respectively. Action potentials were elicited by injecting increasing steps of current (0.5 s) from −25 pA to 250 pA with increments of 25 pA at the cells’ resting potential. The threshold potential was extracted from the first action potential elicited in response to a series of depolarizing current injections and was measured on the ascending phase where the slope exceeded 20 mV/ms. The first current stimulus step that elicited at least one action potential was considered the rheobase (Planert et al., 2023 bioRxiv).

Excitatory postsynaptic currents (EPSCs) in CA1 pyramidal cells were triggered by electrical stimulation of CA3 Schaffer collaterals at increasing current injections ranging from 10 μA - 100 μA for 0.2 ms using Tungsten concentric electrodes (WE3CEA3-200, Science Products) connected to a stimulus isolation unit (WPI A320). CA1 pyramidal cells were voltage clamped at the estimated equilibrium potential for [Cl^-^] at −70 mV to largely suppress inhibitory GABAergic currents.

To determine paired-pulse ratios, half-maximal CA1 EPSCs were evoked by paired stimuli at an inter-stimulus interval of 50 ms. Pulses were triggered at 10 s intervals and 6 individual traces were collected and averaged to determine the EPSC amplitudes and their ratio.

Spontaneous EPSCs (sEPSCs; action potential-dependent and -independent events) and miniature EPSCs (mEPSCs; action potential-independent) were recorded in CA1 pyramidal cells voltage clamped at −70 mV. For mEPSCs, 0.3 μM tetrodotoxin (TTX) was added to the ACSF to block action potentials. Synaptic currents were identified using the template search routine in Clampfit (pClamp10). AMPA current decay was determined using a mono exponential fit.

AMPA/NMDA ratios were determined from mEPSCs by first measuring mixed AMPA- and NMDA receptor-mediated currents in Mg^2+^-free ACSF containing 0.3 μM TTX, 10 μM glycine and 10 μM gabazine, followed by application of 50 μM D-AP5 to isolate the AMPA current component. Average mEPSCs determined from 50 individual events per condition were superimposed and time-matched for current onset to calculate the NMDA receptor component by subtracting the average AMPA mEPSC from the average mixed AMPA+NMDA mEPSC. Individual AMPA and NMDA receptor-mediated components were analyzed as charge entering the cell by calculating respective areas under the mEPSC using the *Area* function in Clampfit (pClamp10).

For single event analysis of mixed AMPA/NMDA synapses, the peak of every mEPSC (AMPA peak) was measured as the mean current over a 1 ms window. NMDA current amplitude (NMDA measurement) was measured over a 5 ms window from 22 to 27 ms after the AMPA peak, a time at which isolated AMPA-mediated mEPSCs had almost completely terminated and declined to 1.08 ± 0.42 pA (n = 15) (Myme et al., 2003). Only events with peak amplitudes of ≥ −30 pA that were significantly larger than the root mean square (RMS) noise of baseline current under Mg^2+^-free conditions were included in this analysis.

To analyze tonic NMDA receptor currents, D-AP5 sensitive RMS noise was measured by averaging holding (baseline) current amplitudes in 10 different regions of 100 ms duration lacking postsynaptic currents, before and after application of D-AP5. D-AP5-sensitive current was obtained by determining the change in mean holding current in a 10 s window before and after D-AP5 application.

### Immunohistochemistry and confocal microscopy

Slices were fixed overnight in 0.1 M phosphate buffered saline (PBS, 0.9% NaCl) containing 4% paraformaldehyde and 4% sucrose at 4°C. After washing with PBS (3 x 15 min), slices were transferred into blocking solution (10% Normal Goat Serum (NGS) and 0.5% Triton-X in PBS) for 1 h. Thereafter, slices were incubated with primary antibody in PBS containing 5% NGS and 0.3% Triton-X for 48-72 h at 4°C. Microglia and astrocytes were labeled using polyclonal rabbit anti-Iba-1 (cat. No. 234 003, 1:1000; Synaptic Systems, Göttingen, Germany) and anti-GFAP antibody (cat. No. 173 308, 1:1000; Synaptic Systems, Göttingen, Germany), respectively. Pre- and post-synapses were labeled using polyclonal guinea pig anti-VGluT1 (cat. No. 135 304, 1:500; Synaptic Systems) and polyclonal rabbit anti-Homer1 (cat. No. 160 002, 1:500; Synaptic Systems). Subsequently, slices were rinsed several times with PBS and then incubated with secondary antibody in PBS containing 3% NGS and 0.1% Triton-X overnight, using Alexa Fluor-488 goat anti-rabbit (1:1000; Jackson, West Grove, USA). Alexa Fluor-488 goat anti-rabbit (A-11008, 1:500 Thermo Fisher) and Alexa Fluor-647 goat anti-guinea pig (A21450, 1:500; Invitrogen) were used for labeling synapses. After washing in PBS, slices were mounted onto 300 μm-thick metal spacers (Bolduan et al., 2020) using aqueous mounting medium Fluoromount-G (Southern Biotech, Birmingham, USA) or Fluoroshield mounting medium with DAPI (Abcam, Cambridge, UK) and stored at 4°C.

Confocal images were collected using Olympus x30 (silicon oil-immersion, 1.05 N.A., 0.8 mm W.D.) and x60 objectives (silicon oil-immersion, 1.35 N.A., 0.15 mm W.D.) on an upright confocal microscope (BX61, Olympus, Tokyo, Japan) with FluoView FV1000 V4.2 acquisition software (Olympus), through separate channels and temporally non-overlapping excitation of the fluorophores, to prevent nonspecific signals. DAPI and Alexa Fluor-647 were excited with diode lasers at 405 nm and 635 nm, respectively, and Alexa Fluor-488 with a multi-line argon laser at 488 nm.

### 3D reconstruction and spine density analysis

Morphology of CA1 pyramidal cell dendrites was analyzed by filling patch-clamped cells with biocytin (0.1%, ≥ 20 min) added to the intracellular solution. After the recordings, slices were fixed and the neurons visualized using Streptavidin-Alexa Fluor-647 conjugate (1:1000; Molecular Probes, Eugene, Oregon, USA) which was included with the secondary antibodies during immuno-labeling as outlined above. High-resolution confocal image stacks using 30x or 60x objectives were taken to visualize pyramidal cell morphology with their apical dendrites and representative fractions of the dendritic shaft and oblique branches. Image stacks were stitched in Fiji and neurons 3D-reconstructed using neuTube software (https://neutracing.com). Per individual neuron, five representative regions of interest comprising 15-20 μm long segments of apical dendritic regions were analyzed. For each individual segment, a 3D skeleton was generated and analyzed for spine number and length using Fiji. Final values for spine density and length per individual cell reflect averages from all five segments.

### Analysis of microglial and astrocyte morphology in brain slices

Slices of every 3D image stack were filtered with a median filter, background subtracted and then transformed into 2D by maximum intensity projection. Microglial coverage was measured through binarization of the images using the Otsu algorithm and the percentage of GFAP area per total area calculated. GFAP intensity was determined as the mean intensity within the GFAP area in 2D-projected images. Microglial morphological parameters were obtained from 3D-Morph automatic image analysis (York et al., 2018). Cell counts were performed in 3D image stacks normalized to a given volume.

### Synapse quantification

Brain slices of 300 μm thickness were cryoprotected in 30 % sucrose/PBS solution and re-sectioned at 70 µm using a cryotome (Microm 340E + Microm KS 34, Thermo Scientific™). The outer slice regions that were most strongly affected by the initial slicing procedure were discarded, and only the inner sections were kept for further use. Image stacks consisting of 12 image planes at 0.25 µm axial (z-axis) intervals were acquired from the CA1 stratum radiatum starting 3 μm below the slice surface using a Leica SP5 confocal microscope with a HCX PL APO lambda blue 63x1.4 Oil objective with additional 3.5x zoom (image field 67.2 µm x 67.2 µm at 1024 x 1024 resolution). Image processing was done using Fiji software. First, images were median filtered (“despeckled”) to remove background noise and maximum intensity projected. Synapses were identified by the colocalization of fluorescent punctae for pre- and postsynaptic markers using the SynapseCounter plugin. This plugin includes image pre-processing (subtraction of smooth continuous background, “Subtract Background”-Plugin; rolling ball radius = 10), binarization and creation of an additive mask for automatic counting of colocalized pre- and postsynaptic elements. Size exclusion (>1.2 μm^2^ and <0.1 μm^2^) was applied to exclude any objects unlikely to represent synapses (McLeod et al., 2017).

### Sholl analysis

CA1 pyramidal cell morphology was assessed by Sholl analysis (Sholl, 1953) using the built in function of the Simple Neurite Tracer plugin in Fiji. Brain slices with biocytin-filled cells were prepared and PFA-fixed as described above. Confocal image stacks of individual CA1 pyramidal cells were acquired at 0.8 µm depth intervals. Image stacks were converted into 8-bit in Fiji and saved as TIF files.

### ELISA-based analysis of basal cytokine and ApoE levels

After removal from the skull, separated brain hemispheres were immediately snap-frozen in liquid nitrogen and kept at −80°C until experimental use. Proteins were extracted from one hemisphere (excluding the cerebellum and olfactory bulb) using a multi-step protocol as described by Kawarabayashi et al. (2001). Briefly, hemispheres were mechanically dissociated with a tissue homogenizer and 1 ml syringe with G26 cannulas. Brain homogenates were sequentially processed in Tris-buffered saline (TBS) followed by Triton-X-containing Tris buffer (TBX) for extraction of soluble (TBS) and membrane-bound proteins (TBX). Proteins dissolved in each buffer were extracted by ultracentrifugation at 100,000 g for 45 min. Supernatants of respective TBS and TBX fractions were collected and stored at −80°C for further analyses. Cytokine and ApoE concentrations of TBS and TBX fractions were determined without further dilution (cytokines) or diluted 1:1000 (ApoE), using V-PLEX Pro-inflammatory Panel 1 (Meso Scale Discovery, catalogue no. K15048D1) and mouse apolipoprotein E ELISA kit (abcam, catalogue no. ab215086) according to the manufacturers’ instructions. Cytokines and ApoE levels were normalized to the total amount of protein determined by BCA protein assay (Thermo Scientific, Pierce, catalogue no. 23225).

### Statistics

Data are presented as means + s.e.m. Statistical tests were performed using OriginPro 9.8 and IBM SPSS statistics (Version: 29.0.0.0 - 241). P values are from unpaired Student’s t-tests (for normally distributed data) or Mann-Whitney U tests (for non-normally distributed data). Normality of data was determined using Anderson-Darling and Shapiro-Wilk tests and equality of variance confirmed using the F-test. Two-way ANOVA followed by Bonferroni post-hoc analysis were used for analysis of the input–output relationship and excitability measurements. For multiple comparisons, p values were corrected using an equivalent to the Holm-Bonferroni method (for N comparisons, the most significant p value is multiplied by N, the 2nd most significant by N-1, the 3rd most significant by N-2, etc.). For analysis of single event mixed AMPA/NMDA mEPSC Pearson correlation and analysis of covariance (ANCOVA) was used. P values were considered significant at ≤ 0.05.

## Acknowledgements

The authors wish to thank Henrik Alle, David Attwell, Jörg Geiger, David Hume and Claire Pridans for comments on the manuscript, and Christian Böttcher, Mehreen Mohammad and Andrea Wilke for expert technical assistance. This work was supported in part by grants from the German Research Foundation (SFB/TRR 167 A10N to C.M. and B07 to J.P.) and the UK DRI (Programme Award to J.P.).

## Expanded View Figure Legends

**Figure EV1:**
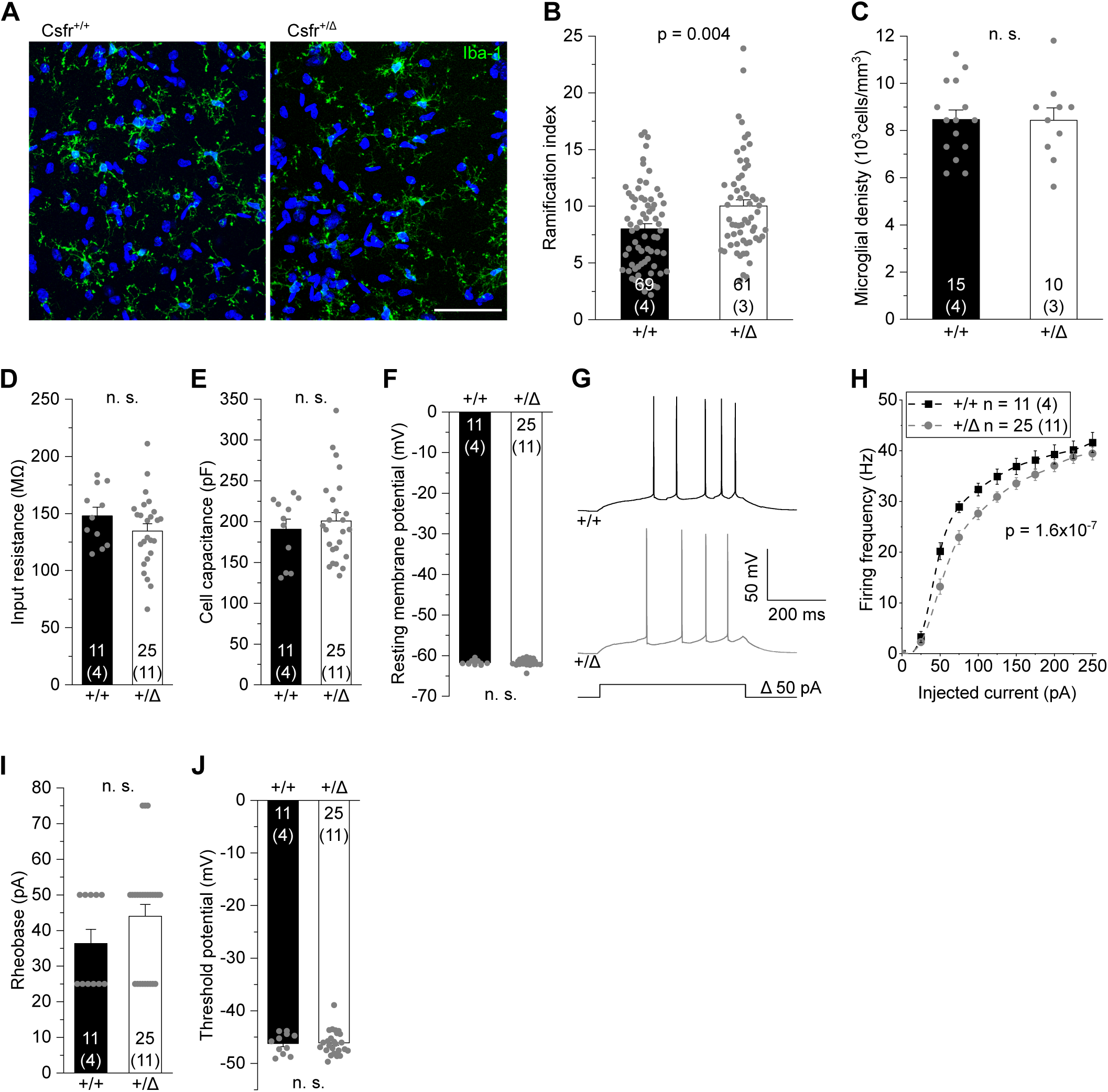
Reduced excitability of CA1 pyramidal cells in heterozygous *Csf1r*^+/ΔFIRE^ mice. A. Specimen confocal images illustrating microglia by Iba1 immunoreactivity (green) in acute hippocampal slices of *Csf1r*^+/ΔFIRE^ (+/Δ) and WT littermates (+/+). DAPI labeling of cellular nuclei in blue. Scale bar, 50 µm B, C Quantification of microglial ramification (B) and cell density (C) in the CA1 *stratum radiatum*. D-F. Analysis of input resistance (D), cell capacitance (E) and resting membrane potential (F) of CA1 pyramidal cells. G. Specimen traces showing action potential firing patterns of CA1 pyramidal cells in response to 500 ms depolarizing current injections. H. Corresponding course of action potential firing frequencies of CA1 pyramidal cells on increasing depolarizations. I, J. Values for rheobase (I) and action potential threshold voltage (J). Data information: Data indicate mean ± SEM. Numbers on bars show tested cells or number of slices (C) and (number of animals). P-values are from unpaired Student’s t (B – F, J) or Mann-Whitney tests (I) and two-way ANOVA (H).

**Figure EV2:**
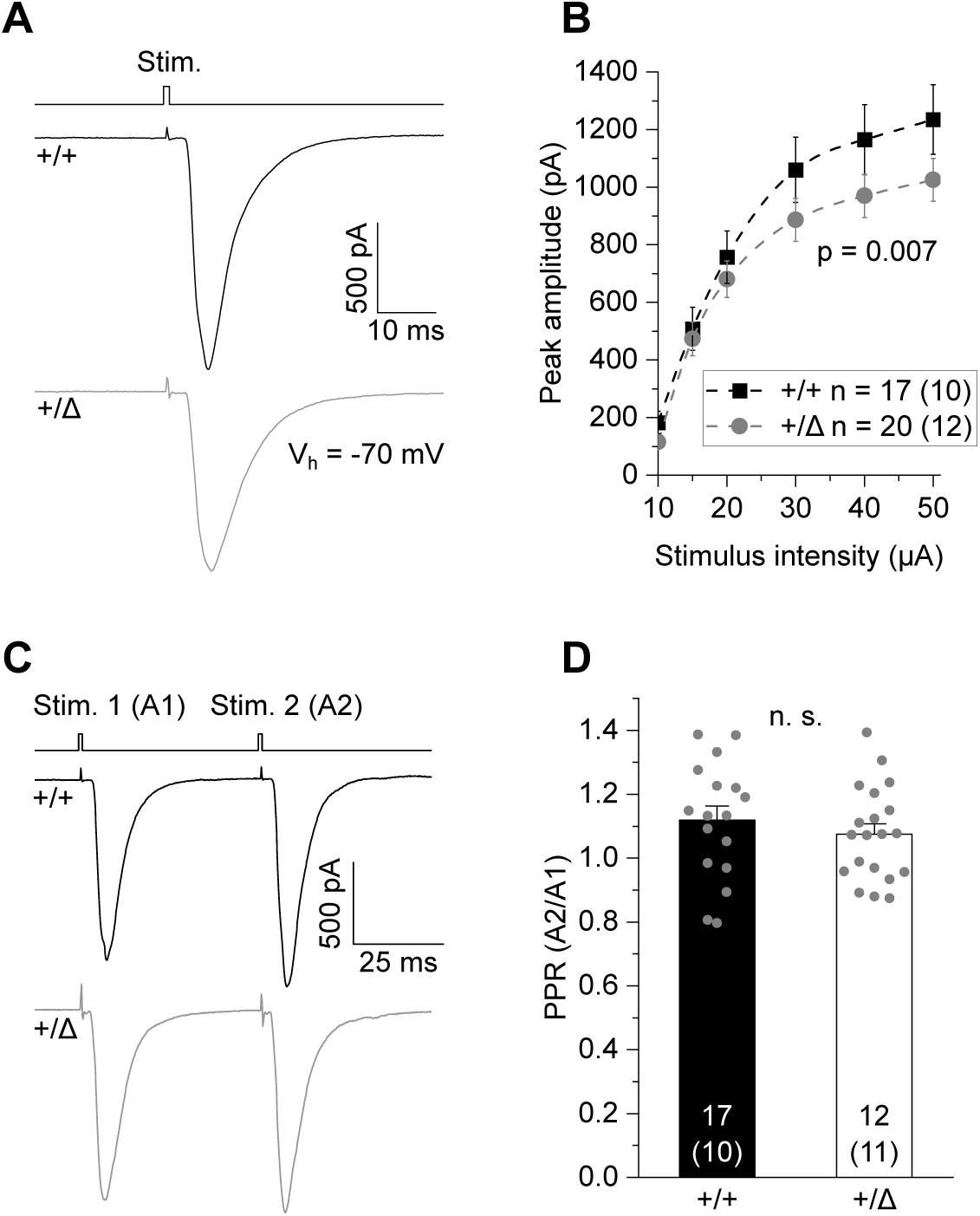
Reduced CA3 – CA1 glutamatergic transmission in heterozygous *Csf1r*^+/ΔFIRE^ mice. A. Specimen traces showing excitatory postsynaptic currents (EPSC) in CA1 pyramidal cells in response to electrical stimulation for 0.2 ms in +/Δ and +/+ mice. B. Corresponding input-output relationships showing changes in peak amplitudes of CA1 EPSCs with increasing stimulation strength. C. Example traces (EPSCs) of CA1 pyramidal cells after paired-pulse stimulation at 50 ms inter-stimulus intervals. D. Comparison of paired pulse ratios (PPR) as the quotient of the second vs first EPSC amplitude (A2/A1) of CA1 pyramidal cells. Data information: Data are represented as mean ± SEM. Numbers on bars indicate tested cells and (number of animals). P-values are from two-way ANOVA (B) and unpaired Student’s t test (D).

**Figure EV3:**
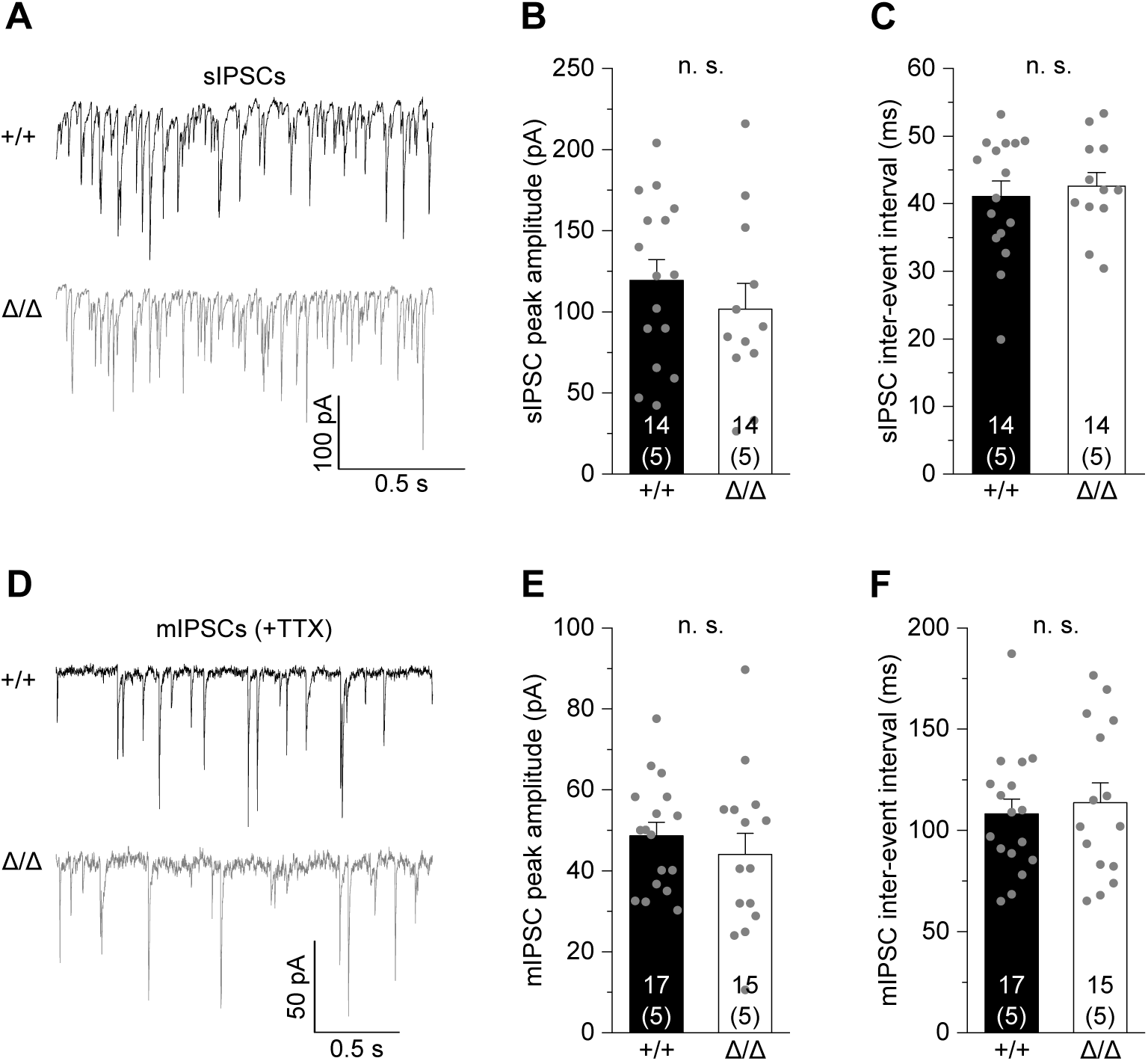
Unaltered function of inhibitory synapses of CA1 pyramidal cells in Csf1r^ΔFIRE/ΔFIRE^ mice. A. Specimen traces showing GABA_A_ receptor-evoked spontaneous inhibitory post-synaptic currents (sIPSCs) of CA1 pyramidal cells in +/+ and Δ/Δ mice. B, C. Comparison of sIPSC peak amplitudes (B) and inter-event intervals (C). D. Specimen traces showing miniature IPSCs (mIPSCs) of CA1 pyramidal cells. E, F. Comparison of peak amplitudes (E) and inter-event intervals (F). Data information: Data indicate mean ± SEM. Numbers on bars show tested cells and (number of animals). P-values are from unpaired Student’s t tests.

**Figure EV4:**
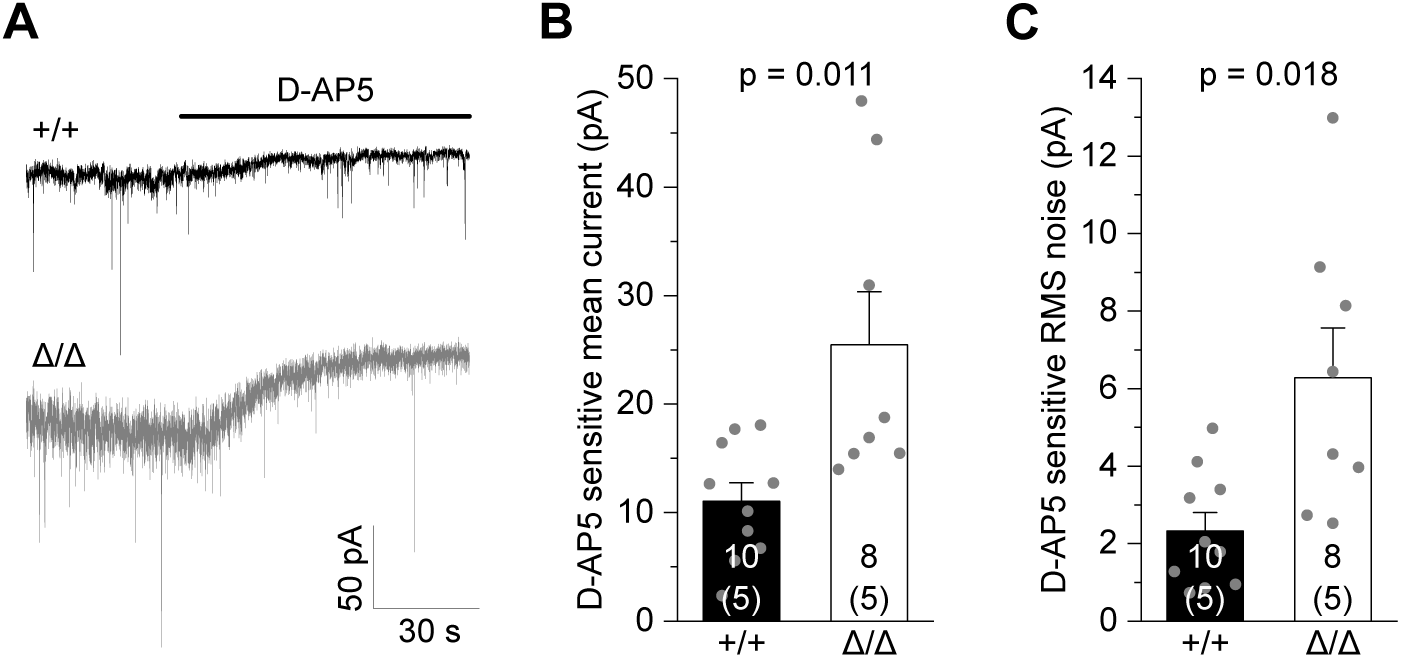
Increased tonic NMDA current in *Csf1r*^ΔFIRE/ΔFIRE^ mice. A. Example traces showing changes in holding current after application of 50 µM D-AP5, reflecting blockade of all tonic NMDAR-mediated currents in CA1 pyramidal cells. Measurements were done in nominally Mg^2+^-free extracellular solution in the presence of 300 nM TTX and 10 μM glycine. B, C. Comparison of the D-AP5-sensitive mean holding current (B) and root mean square (RMS) noise (C) between genotypes. Note the contribution of both synaptic and extrasynaptic NMDA receptors. Data information: Data are represented as mean ± SEM. Numbers on bars indicate tested cells and (number of animals). P-values are from Mann-Whitney (B) or unpaired Student’s t tests (C).

